# Probabilistic Learning of Treatment Trees in Cancer

**DOI:** 10.1101/2022.01.23.477414

**Authors:** Tsung-Hung Yao, Zhenke Wu, Karthik Bharath, Jinju Li, Veerabhadran Baladandayuthapan

## Abstract

Accurate identification of synergistic treatment combinations and their underlying biological mechnisms is critical across many disease domains, especially cancer. In translational oncology research, preclinical systems such as patient-derived xenografts (PDX) have emerged as a unique study design evaluating multiple treatments administered to samples from the same human tumor implanted into genetically identical mice. In this paper, we propose a novel Bayesian probabilistic tree-based framework for PDX data to investigate the hierarchical relationships between treatments by inferring treatment cluster trees, referred to as treatment trees (R_x_-tree). The framework motivates a new metric of mechanistic similarity between two or more treatments accounting for inherent uncertainty in tree estimation; treatments with a high estimated similarity have potentially high mechanistic synergy. Building upon Dirichlet Diffusion Trees, we derive a closed-form marginal likelihood encoding the tree structure, which facilitates computationally efficient posterior inference via a new two-stage algorithm. Simulation studies demonstrate superior performance of the proposed method in recovering the tree structure and treatment similarities. Our analyses of a recently collated PDX dataset produce treatment similarity estimates that show a high degree of concordance with known biological mechanisms across treatments in five different cancers. More importantly, we uncover new and potentially effective combination therapies that confer synergistic regulation of specific downstream biological pathways for future clinical investigations. Our accompanying code, data, and shiny application for visualization of results are available at: https://github.com/bayesrx/RxTree.

## 1 Introduction

According to the World Health Organization, cancer is one of the leading causes of death globally, with ∼10 million deaths in 2020 [Ferlay et al., 2020]. Despite multiple advances over the years, systematic efforts to predict efficacy of cancer treatments have been stymied due to multiple factors, including patient-specific heterogeneity and treatment resistance [Dagogo-Jack and Shaw, 2018, Groisberg and Subbiah, 2021]. Given that the evolution of tumors relies on a limited number of biological mechanisms, there has been a recent push towards combining multiple therapeutic agents, referred to as “combination therapy” [Sawyers, 2013, Groisberg and Subbiah, 2021]. This is driven by the core hypothesis that combinations of drugs act in synergistic manner, with each drug compensating for the drawbacks of other drugs. However, despite higher response rates and efficacy in certain instances [Bayat Mokhtari et al., 2017], combination therapy can lead to undesired drug-drug interactions, lower efficacy, or severe side effects [Sun et al., 2016]. Consequently, it is highly desirable to advance the understanding of underlying mechanisms that confer synergistic drug effects and identify potential favorable drug-drug interaction mechanisms for further investigations.

Given that not all possible drug combinations can be tested on patients in actual clinical trials, cancer researchers rely on preclinical “model” systems to guide the discovery of the most effective combination therapies (note, models have a different contextual meaning here). In translational oncology, preclinical models assess promising treatments and compounds, before they are phased into human clinical trials. The traditional mainstay of such preclinical models has been cell-lines, wherein cell cultures derived from human tumors are grown in an *in vitro* controlled environment. However, it has been argued that they do not accurately reflect the true behavior of the host tumor and, in the process of adapting to *in vitro* growth, lose the original properties of the host tumor, thus leading to limited clinical relevance and successes [Tentler et al., 2012, Bhimani et al., 2020]. To overcome these challenges, there has been a push towards more clinically relevant model systems that maintain a high degree of fidelity to human tumors. One such preclinical model system is Patient-Derived Xenograft (PDX) wherein tumor fragments obtained from cancer patients are directly transplanted into genetically identical mice [Hidalgo et al., 2014, Lai et al., 2017]. Compared to traditional oncology models such as cell-lines [Yoshida, 2020], PDX models maintain key cellular and molecular characteristics, and are thus more likely to mimic human tumors and facilitate precision medicine. More importantly, accumulating evidence suggests responses (e.g. drug sensitivity) to standard therapeutic regimens in PDXs closely correlate with patient clinical data, making PDX an effective and predictive experimental model across multiple cancers [Topp et al., 2014, Nunes et al., 2015].

### PDX experimental design and key scientific questions

Briefly, in a standard PDX preclinical experiment, a set of common treatments are tested, and each treatment is given to multiple mice with tumors implanted from the same (matched) patient (see conceptual schema in Figure 1(A)). Treatment responses (e.g. tumor size) are then evaluated, resulting in a data matrix (treatments × patients) as depicted in the heatmap in Figure 1(A). The PDX experimental protocol, often referred to as “co-clinical trials,” mirrors a real human clinical trial using mouse “avatars” [Clohessy and Pandolfi, 2015]. Thus the protocol serves as a scalable platform to: (a) identify underlying plausible biological mechanisms responsible for tumor growth and resistance, and (b) evaluate the effectiveness of drug combinations based on mechanistic understanding [Lunardi and Pandolfi, 2015]. In this context, the (biological) mechanism refers to the specific mechanism of action of a treatment, which usually represents a specific target, such as an enzyme or a receptor [Grant et al., 2010]. From the perspective of treatment responses as data, responses are the consequences of the downstream biological pathways from the corresponding interaction between a treatment and the target/mechanism.

**Figure 1:**
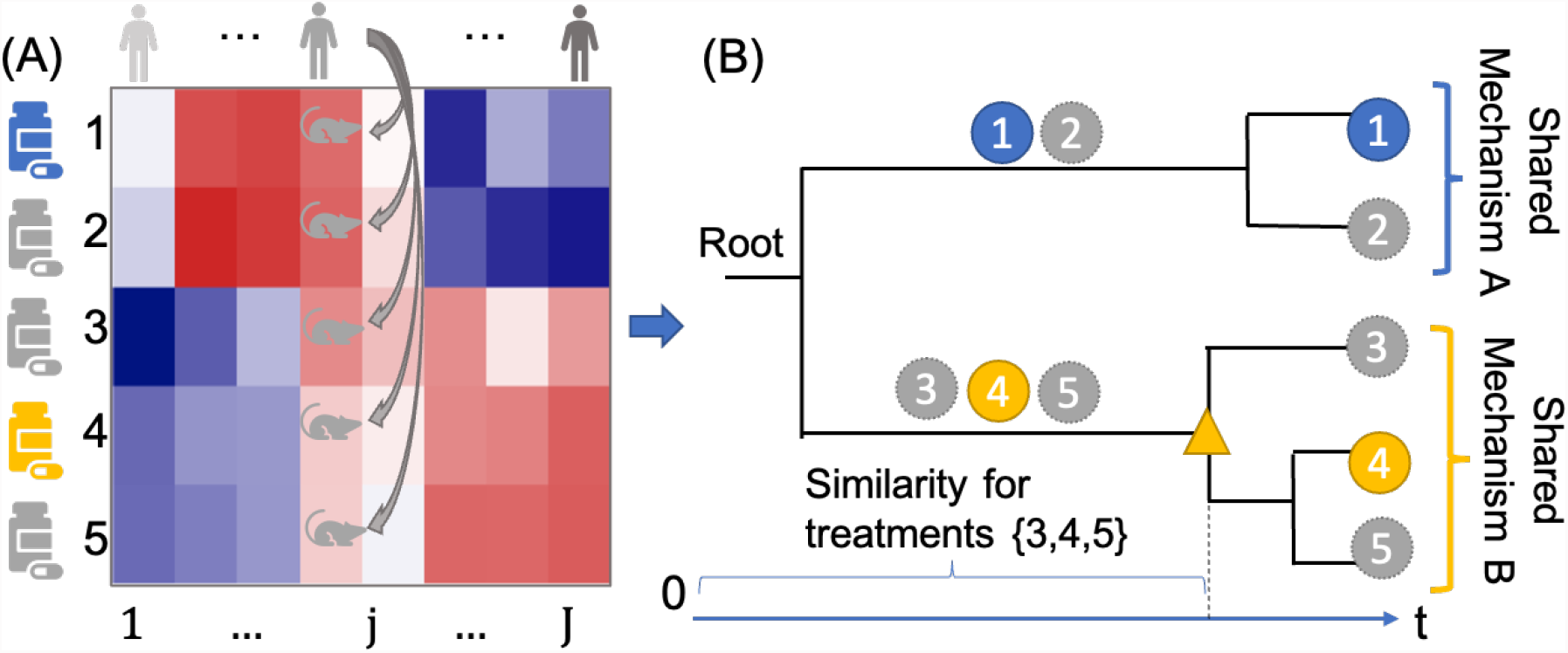
PDX experimental design and tree-based representation. Panel **A**: an illustrative PDX dataset with five treatments (row) and eight patients (column). Mice in a given column are implanted with tumors from the same patient and receive different treatments (across rows). The level of tumor responses are shown along a color gradient. Panel **B**: a tree structure that clusters the treatments and quantifies the similarity among mechanisms. Two treatments (1 and 4) are assumed to have different but known biological mechanisms (in different colors); the rest three treatments (2,3, and 5) have unknown mechanisms (in gray). The tree suggests two treatment groups are present ({1, 2} and {3, 4, 5}) that may correspond to two different known mechanisms. The horizontal position of “△” represents the divergence time (defined in Section 2.1) and the mechanism similarity for treatments {3, 4, 5 }. In a real data analysis, the tree (topology and divergence times) is unknown and is to be inferred from PDX data.

Ideally, treatments with the same target/mechanism should induce similar responses and engender mechanism-related clustering among treatments. Evidently then, a sensible clustering of treatments would not only partition treatments into clusters but also explicate how the clusters relate to one another; in other words, a hierarchy among treatment clusters is more likely to uncover plausible mechanisms for combinations of treatments with “similar” responses when compared to “flat” clusters (e.g., *k*-means clustering). Such response-based identification of potential synergistic effects from combinations of treatments will augment understanding from known mechanistic synergy. In our application, using tree-based clustering, we assume known entities at the leaves, i.e., the different treatments. The treatments are assumed to act upon potentially distinct biological pathways, resulting in different levels of responses across the treated mice. In this paper, we use PDX response data on the leaves to infer a hierarchy over treatments that may empirically characterize the similarity in the targeted mechanistic pathways. The primary statistical goals are to (i) define and estimate a general metric measuring the similarity within any subset comprising two or more treatments, and (ii) facilitate (i) by conceptualizing and inferring an unknown hierarchy among treatments.

### Tree-based representations for PDX data

To this end, we consider a tree-based construct to explore the hierarchical relationships between treatments, referred to as *treatment tree (R*_*x*_*-tree*, in short). We view such a tree structure as a representation of clustering of treatments based on mechanisms that confer synergistic effects, wherein similarities between mechanisms are captured through branch lengths. Hierarchy among treatments can be interpreted through branch lengths (from the root) that are potentially reflective of different cancer processes; this would then help identify common mechanisms and point towards treatment combinations disrupting oncological processes if administered simultaneously.

We will focus on rooted trees. The principal ingredients of a rooted tree comprise a root node, terminal nodes (or, leaves), internal nodes and branch lengths. In the context of the R_x_-tree for PDX data, the leaves are observed treatment responses, whereas internal nodes and branch lengths are unobserved. Internal nodes are clusters of treatments, and lengths of branches between nodes are indicative of strengths of mechanism similarities. The root is a single cluster consisting of all treatments. This leads to the following interpretation: at the root all treatments share a common target or mechanism; length of path from the root to the internal node (sum of branch lengths) at which two treatments split into different clusters measures mechanism similarity between the two treatments. Thus treatments that stay clustered “longer” have higher mechanism similarities.

Throughout, we will use ‘tree’ when describing methodology for an abstract tree (acyclic graphs with distinguished root node) and ‘treatment tree’ or ‘R_x_-tree’ when referring to the latent tree within the application context.

### An illustrative example

A conceptual R_x_-tree and its interpretation is illustrated in Figure 1 where five treatments (1 to 5) are applied on eight patients’ PDXs (Figure 1(A)) with the corresponding (unknown true) R_x_-tree (Figure 1(B)) based on the PDX data. Assume two treatment groups based on different mechanisms – treatments {1, 2} and treatments {3, 4, 5 }; further, suppose treatment 4 is approved by the Food and Drug Administration (FDA). The heatmap in Panel (A) visualizes the distinct levels of response profiles to the five treatments so that treatments closer in the tree are more likely to have similar levels of responses. The R_x_-tree captures the mechanism similarity by arranging treatments {1, 2}and {3, 4, 5} to stay in their respective subtrees longer and to separate the two sets of treatment early in the tree. Based on the R_x_-tree, treatments {3, 5} share high mechanism similarity values with treatment 4; treatment 5 is the closest to the treatment 4, suggesting the most similar synergistic mechanism among all the evaluated treatments 1 to 5.

### Existing methods and modeling background

The Pearson correlation is a popular choice to assess mechanism similarity between treatments [Krumbach et al., 2011], but is inappropriate to examine multi-way similarity. A tree-structured approach based on a (binary) dendrogram obtained from hierarchical clustering of cell-line data using the cophenetic distance [Sokal and Rohlf, 1962] was adopted in Narayan et al. [2020]; their approach, however, failed to account for uncertainty in the dendrogram, which is highly sensitive to measurement error in the response variables as well distance metrics (we show this via simulations and in real data analyses). In this paper, we consider a model for PDX data parameterized by a tree-structured object representing the R_x_-tree. The model is derived from the Dirichlet diffusion tree (DDT) [Neal, 2003] generative model for (hierarchically) clustered data. The DDT engenders a data likelihood and a prior distribution on the tree parameter with support in the space of rooted binary trees. We can then use the posterior distribution to quantify uncertainty about the latent R_x_-tree.

### Summary of novel contributions and organization of the article

Our approach based on the DDT model for PDX data results in three main novel contributions:

a. *Derivation of a closed-form likelihood that encodes the tree structure.* The DDT specification results in a joint distribution on PDX data, treatment tree parameters and other model parameters. By marginalizing over unobserved data that correspond to internal nodes of the tree, we obtain a new multivariate Gaussian likelihood with a special tree-structured covariance matrix, which completely characterizes the treatment tree (Proposition 1 and Lemma 1).
b. *Efficient two-stage algorithm for posterior sampling.* Motivated by the form of marginal data likelihood in (a), we decouple the Euclidean and tree parameters and propose a two-stage algorithm that combines an approximate Bayesian computation (ABC) procedure (for Euclidean parameters) with a Metropolis-Hasting (MH) step (for tree parameters). We demonstrate via multiple simulation studies the superiority of our hybrid approach over approaches based on classical single-stage MH algorithms (Sections 4.2 and 4.1).
c. *Corroborating existing, and uncovering new, synergistic combination therapies.* We define and infer a new similarity measure that accounts for inherent uncertainty in estimating a latent hierarchy among treatments. As a result, the *maximum a posteriori* R_x_-tree and the related mechanism similarity show high concordance with known existing biological mechanisms for monotherapies and uncover new and potentially useful combination therapies (Sections 5.3 and 5.4).

Of particular note is contribution (c), where we leverage a We calculated the Novartis Institutes for <u>B</u>ioMedical <u>R</u>esearch - <u>PDX E</u>ncyclopedia [NIBR-PDXE, [Gao et al., 2015]] that interrogated multiple targeted therapies across five different cancers. Our pan-cancer analyses of the NIBR-PDXE dataset show a high degree of concordance with known existing biological mechanisms across different cancers; for example, a high mechanistic similarity is suggested between two agents currently in clinical trials: CGM097 and HDM201 in breast cancer and colorectal cancer, known to target the same gene MDM2 [Konopleva et al., 2020]. In addition, our model uncovers new and potentially effective combination therapies. For example, exploiting knowledge of the combination therapy of a class of agents targeting the PI3K-MAPK-CDK pathway axes – PI3K-CDK for breast cancer, PI3K-ERBB3 for colorectal cancer and BRAF-PI3K for melanoma – confers possible synergistic regulation for prioritization in future clinical studies.

The rest of the paper is organized as follows: we first review our probabilistic formulation for PDX data based on the DDT model and present the marginal data likelihood and computational implications in Section 2. In Section 3, we derive the posterior inference algorithm based on a two-stage algorithm. In Section 4, we conduct two sets of simulations to evaluate the operating characteristics of the model and algorithm. A detailed analysis of the NIBR-PDXE dataset, results, biological interpretations and implications are summarized in Section 5. The paper concludes by discussing implications of the findings, limitations, and future directions.

## 2 Modeling R_x_-tree via Dirichlet Diffusion Trees

Given a PDX experiment with *I* correlated treatments and *J* independent patients, we focus on the setting with 1× 1 ×1 design (one animal per PDX model per treatment) with no replicate response for each treatment and patient. A PDX experiment produces an observed data matrix **X**_*I×J*_ = [***X***_1_, …, ***X***_*I*_]^T^ where ***X***_*i*_ = [*X*_*i*1_, …, *X*_*iJ*_]^T^ is data under treatment *i* across *J* patients; let the observed response column for each patient be ***X***_.,*j*_ = [*x*_1*j*_, …, *x*_*Ij*_]^T^ ∈ ℝ^*I*^, *j* = 1, …, *J*.

In this paper, the observed treatment responses are continuous and we model the responses through a generative model that results in a Gaussian likelihood with a structured covariance:

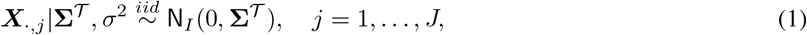

where the **Σ**^𝒯^ is a tree-structured covariance matrix that encodes the tree 𝒯. In particular, 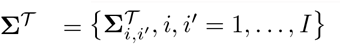 encodes the tree 𝒯 through two constraints [Lapointe and Legendre, 1991, McCullagh, 2006]:

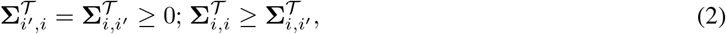

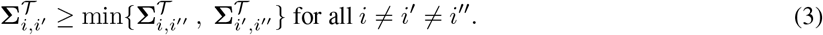

Each element 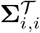 is the covariance between treatments *i* and *i*′ and measures their similarity. The inequality (2) imposes the symmetry of covariance matrix and ensures the divergence of all leaves. The tree structure is characterized by the ultrametric inequality (3) that ensures **Σ**^𝒯^ bijectively maps to a tree 𝒯; for more details on the relationship between the covariance **Σ**^𝒯^ and the tree 𝒯 see McCullagh [2006] and Bravo et al. [2009]. Of note, mean parameterized models (e.g. mixed effects models) are inappropriate for uncovering the tree parameter under the given data structure since the latent tree is completely encoded in covariance matrix **Σ**^𝒯^

A Bayesian formulation requires an explicit prior distribution on **Σ**^𝒯^ which satisfies constrains (2) and (3); this requirement is far from straightforward since the set of tree-structured matrices is complicated (e.g., it is not a manifold [McCullagh, 2006]). We instead consider the Dirichlet Diffusion tree (DDT) model [Neal, 2003] for hierarchically clustered data which provides two useful ingredients:

1. a prior is implicitly specified on the latent treatment tree, comprising the root, internal nodes, leaves, and branch lengths;
2. upon integrating out the internal nodes, a tractable Gaussian likelihood on PDX data with tree-structured covariance is specified.

We first provide a brief description of the DDT model proposed by Neal [2003] and its joint density on data and tree (Section 2.1). Subsequently, we derive an expression for the likelihood and demonstrate how it can be profitably employed to develop a generative model for PDX data and carry out R_x_-tree estimation (Section 2.2 and 2.3).

### 2.1 The Generative Process of DDT

The DDT prescribes a fragmentary, top-down mechanism to generate a binary tree (acyclic graph with a preferred node or vertex referred to as the root), starting from a root, containing *J* -dimensional observed responses ***X***_*i*_ at *I* leaves/terminal nodes; each node in the tree has either 0 or 2 children excepting the root which has a solitary child. This prescription manipulates dynamics of a system of *I* independent Brownian motions *B*_1_, …, *B*_*I*_ on ℝ^*J*^ in a common time interval *t* ∈ [0, 1]. As shown in Figure 2(A), all Brownian motions *B*_*i*_(*t*) start at the same point at time *t* = 0, location of which is the root **0** ∈ ℝ^*J*^, and diverge at time points in [0, 1] and locations in ℝ^*J*^ before stopping at the time *t* = 1 at locations ***X***_*i*_. The Brownian trajectories and their divergences engender the tree structure as shown in Figure 2(A).

**Figure 2:**
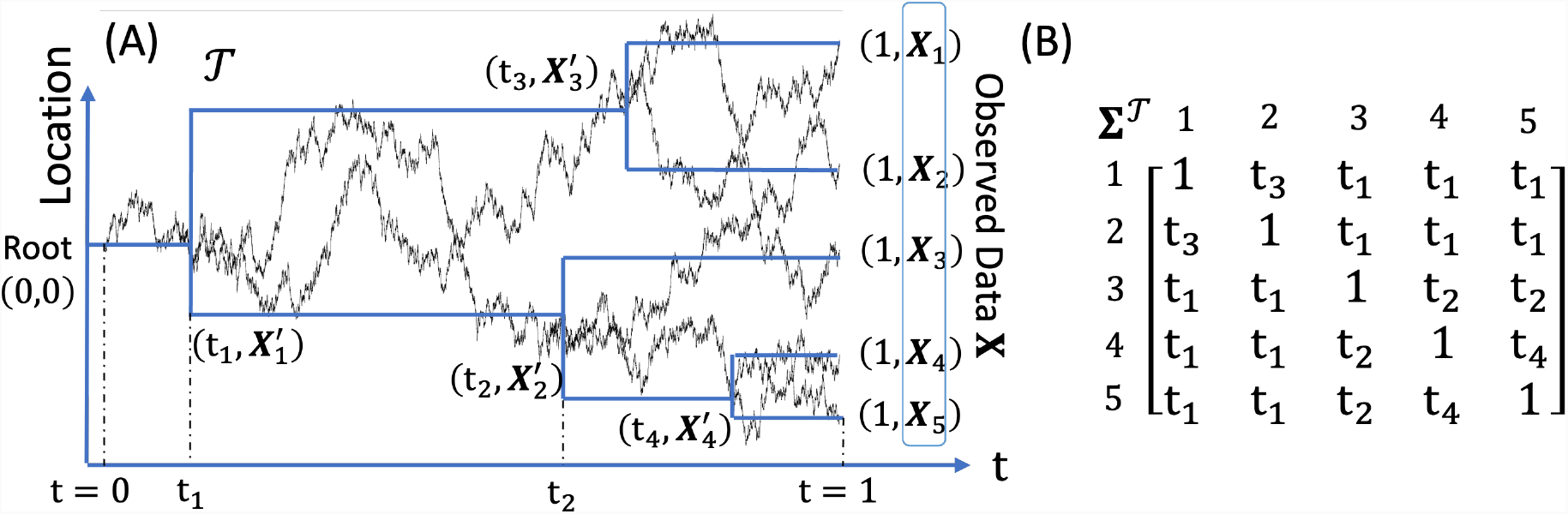
(A) A binary tree with *I* = 5 leaves underlying the diffusion dynamics. The observed response vector ***X***_*i*_, *i* = 1, …, *I* is generated by the Brownian motion up to *t* = 1. The unobserved response vector 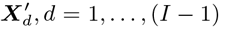 at the divergence is generated by the Brownian motion at time *t*_*d*_. (B) A tree-structured matrix **Σ**^𝒯^ that encapsulates the tree 𝒯. See the Proposition 1 for the definition of **Σ**^𝒯^.

Specifics on when and how the Brownian motions diverge are as follows: the first Brownian motion *B*_1_(*t*) starts at *t* = 0 and generates ***X***_1_ at *t* = 1; a second independent Brownian motion *B*_2_(*t*) starts at the same point at *t* = 0, branches out from the first Brownian motion at some time *t*, after which it generates ***X***_2_ at time 1. The probability of divergence in a small interval [*t, t* + *dt*] is given by a *divergence function t* ↦ *a*(*t*), assumed as in Neal [2003] to be of the form *a*(*t*) = *c*(1 − *t*)^−1^ for some divergence parameter *c >* 0. Inductively then, the vector of observed responses to treatment *i*, ***X***_*i*_, is generated by *B*_*i*_(*t*), which follows the path of previous ones. If at time *t, B*_*i*_(*t*) has not diverged and meets the previous divergent point, it will follow one of the existing path with the probability proportional to the number of data points that have previously traversed along each path. Eventually, given *B*_*i*_(*t*) has not diverged at time *t*, it will do so in [*t, t* + *dt*] with probability *a*(*t*)*dt/m*, where *m* is the number of data points that have previously traversed the current path.

From the illustration in panel (A) of Figure 2, we note that *B*_3_ diverges from the *B*_1_ and *B*_2_ at time *t*_1_ at location 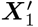 and at *t* = 1 is at location ***X***_3_, which is the *J*-dimensional response vector for treatment 3; this creates a solitary branch of length *t*_1_ from the root and an unobserved internal node at location 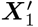. Continuing, given three Brownian motions *B*_1_, *B*_2_ and *B*_3_, *B*_4_ does not diverge before *t*_1_ and meet the previous divergent point *t*_1_. *B*_4_ chooses to follow the path of *B*_3_ with probability 1*/*3 at *t*_1_ and finally diverges from *B*_3_ at time *t*_2_ *> t*_1_ at location 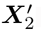; this results in observation ***X***_4_ for treatment 4 and an unobserved internal node at 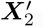, and so on. As a consequence, the binary tree that arises from the DDT comprises of:

i. an unobserved root at the origin in ℝ^*J*^ at time *t* = 0;
ii. observed data **X** = [***X***_1_, …, ***X***_*I*_]^T^ ∈ ℝ^*I×J*^ situated at the leaves of the tree;
iii. unobserved internal nodes 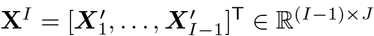;
iv. unobserved times ***t*** = (*t*_1_, …, *t*_*I*−1_)^T^ ∈ [0, 1]^*I*−1^ that characterize lengths of branches;
v. unobserved topology 𝒯 that links (i)-(iv) into a tree structure, determined by the number of data points ***X***_*i*_ that have traversed through each segment or branch.

Conceptually, observed data at the leaves ***X***_1_, …, ***X***_*I*_ collectively form the observed PDX responses generated through a process involving a few parameters: tree-related parameters (𝒯, ***t***) and the locations of internal nodes 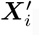 The tree 𝒯 clusters *I* treatments as a hierarchy of (*I* − 1) levels (excluding the last level containing leaves). At level 0 *< d* ≤ *I* − 1 of the hierarchy, characterized by the pair 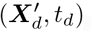, the *I* treatments are clustered into *d* + 1 groups; a measure of similarity (or dissimilarity) between treatment clusters at levels *d* and *d* + 1 is given by the branch length *t*_*d*+1_ − *t*_*d*_.

We now give a brief description of how the joint density of (**X, X**^*I*^, ***t***, *𝒯*) can be derived; for more details we direct the reader to Neal [2003] and Knowles and Ghahramani [2015]. For a fixed *c >* 0 that governs the divergence function *a*(*t*) = *c*(1− *t*)^−1^, probabilities associated with the independent Brownian motions *B*_1_, …, *B*_*I*_ induce a joint (Lebesgue) density on the generated tree. Note that the binary tree arising from the DDT is encoded by the triples 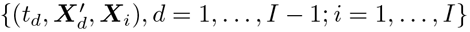. An internal node at 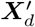 contains *l*_*d*_ and *r*_*d*_ leaves below to its left and right with *m*_*d*_ = *l*_*d*_ + *r*_*d*_. If each of the Brownian motions is scaled by *σ*^2^ *>* 0, then given 𝒯 and a branch with endpoints 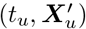 and 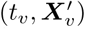 with 0 *< t*_*u*_ *< t*_*v*_ *<* 1, from properties of a Brownian motion we see that 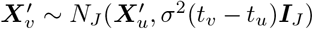, and the (Lebesgue) density of 𝒯 can be expressed as the product of contributions from its branches. Then the joint density of all nodes, times and the tree topology is given by

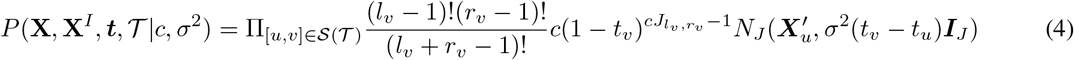

where 𝒮 (𝒯) is the collection of branches and 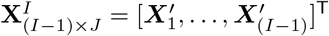 are unobserved locations of the internal nodes. On each branch [*u, v*], the first term 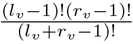 represents the chance the branch containing *l*_*v*_ and *r*_*v*_ leaves to its left and right respectively; 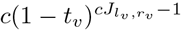 represents the probability of diverging at *t*_*v*_ with *l*_*v*_ and *r*_*v*_ leaves, where 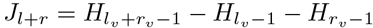 with 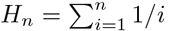 is the *n*th harmonic number.

The joint density is hence parameterized by (*c, σ*^2^), where *c* plays a crucial role in determining the topology 𝒯: through the divergence function *a*(*t*), it determines the propensity of the Brownian motion to diverge from its predecessors; consequently, a small *c* engenders later divergence and a higher degree of similarity among treatments in PDX. The latent tree has two components: (i) topology 𝒯 and (ii) vector of divergence times ***t*** determining branch lengths. We refer to (*c, σ*^2^) as the *Euclidean parameters* and (𝒯, ***t***) as *tree parameters*.

### 2.2 Prior on tree and closed-form likelihood

The joint density in (4) factors into a prior *P* (***t***, 𝒯 |*c, σ*^2^) on the tree parameter through (𝒯, ***t***) and a density *P* (**X, X**^*I*^ |***t***, 𝒯, *c, σ*^2^) that is a product of *J* -dimensional Gaussians on the internal nodes and leaves. The prior distribution on the latent tree is thus implicitly defined through the Brownian dynamics and is parameterized by (𝒯, ***t***) with hyperparameters (*c, σ*^2^). In (4) the product is over the set of branches 𝒮 (𝒯), and the contribution to the prior *P* (𝒯, ***t*** |*c, σ*^2^) from each branch [*u, v*] is 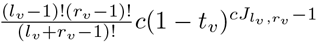, which is free of *σ*^2^; on the other hand, the contribution to *P* (**X, X**^*I*^ |***t***, *𝒯, c, σ*^2^) from [*u, v*] is the *J*-dimensional 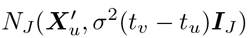, which is independent of *c*. The likelihood function based on the observed **X** is thus obtained by integrating out the unobserved internal nodes **X**^*I*^ from *P* (**X, X**^*I*^ |***t***, *𝒯, σ*^2^). Accordingly, our first contribution is to derive a closed-form likelihood function for efficient posterior computations; to our knowledge, this task is currently achieved only through sampling-based or variational methods [Neal, 2003, Knowles and Ghahramani, 2015].

Denote as `MN`_*I* × *J*_ (*M, U, V*) the matrix normal distribution of an *I* × *J* random matrix with mean matrix *M*, row covariance *U*, and column covariance *V*, and let ***I***_*k*_ denote the *k* × *k* identity matrix. Evidently, **X** follows a matrix normal distribution since Gaussian laws of the Brownian motions imply that 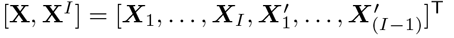 follow a matrix normal distribution.

#### Proposition 1.

Under the assumption that the root is located at the origin in ℝ^*J*^, the data likelihood **X**|*σ*^2^, 𝒯, ***t*** ∼ `MN`_*I×J*_ (**0**, *σ*^2^**Σ**^𝒯^, ***I***_*J*_), where 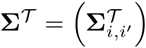 is an *I* × *I* tree-structured covariance matrix satisfying (2) and (3) with 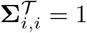and 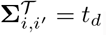, for *i* ≠ *i*′ where *i, i*′ = 1, …, *I* and *d* = 1, …, *I* − 1.

Proposition 1 asserts that use of the DDT model leads to a centered Gaussian likelihood on PDX data ***X*** with a tree-structured covariance matrix. Proposition 1 also implies that each patient independently follows the normal distribution of (1) with an additional scale parameter (*σ*^2^) from the Brownian motion:

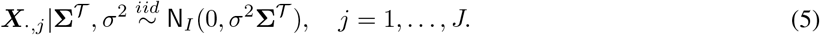

By setting 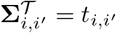 as the divergence time of *i* and *i*′, **Σ**^𝒯^ satisfies (2) and (3) and encodes the tree 𝒯. For example, consider a three-leaf tree with 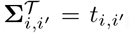, inequality (3) implies that for the three leaves, say, *i, i*′ and *i*″, one of the following conditions must hold: (i) *t*_*i*′,*i*″_ ≥*t*_*i,i*′_ = *t*_*i,i*″_; (ii) *t*_*i*,″_ ≥*t*_*i,i*′_ = *t*_*i*′,*i*″_ ; (iii) *t*_*i,i*′_≥ *t*_*i,i*″_ = *t*_*i*′,*i*″_. We then obtain a tree containing 1) a subtree of two leaves with a higher similarity and 2) a singleton clade with a lower similarity between the singleton leaf and the two leaves in the first subtree. In particular, if *t*_*i*′,*i*″_ ≥*t*_*i,i*′_ = *t*_*i,i*″_ holds, the three-leaf tree has leaf *i* diverging earlier before the subtree of (*i*′, *i*″).

### 2.3 Decoupling Tree and Euclidean Parameters for Efficient Sampling

In the full joint density in (4) the Euclidean and tree parameters are confounded across row and column dimensions of **X**, and this may result in slow mixing of chains using traditional MCMC algorithms [Turner et al., 2013]. State-of-the-art posterior inference on (*c, σ*^2^, *𝒯*, ***t***) can be broadly classified into sampling-based approaches [e.g., Knowles and Ghahramani, 2015] and deterministic approaches based on variational message passing [e.g., Knowles et al., 2011, VMP]. Variational algorithms can introduce approximation errors to the joint posterior via factorization assumptions (e.g.,mean-field) and choice of algorithm is typically determined by the speed-accuracy trade-off tailored for particular applications. On the other hand, in classical MCMC-based algorithms for DDT we observed slow convergence in the sampling chains for *c* and *σ*^2^ with high autocorrelations for the corresponding chains, owing to possibly the high mutual dependence between *c* in the divergence function and the tree topology 𝒯, resulting in slow local movements in the joint parameter space of model and tree parameters (Simulation II in Section 4.2).

Notwithstanding absence of the parameter *c* in the Gaussian likelihood, the dependence, and information about, *c* is implicit: the distribution of divergence times ***t*** that populate **Σ**^𝒯^ are completely determined by the divergence function *t* ↦*c*(1− *t*)^−1^. In other words, *c* can indeed be estimated from treatment responses {***X***_*·,j*_ } using the likelihood. From a sampling perspective, however, form of the likelihood obtained by integrating out the internal nodes **X**^*I*^, suggests an efficient two-stage sampling strategy that resembles the classical collapsed sampling [Liu, 1994] strategy in MCMC literature: first draw posterior samples of (*c, σ*^2^) and then proceed to draw posterior samples of (𝒯, ***t***) conditioned on each sample of (*c, σ*^2^).

## 3 R_x_-tree Estimation and Posterior Inference

In line with the preceding discussion, we consider a two-stage sampler for Euclidean and tree parameters. While in principle MCMC techniques could be used in both stages, we propose to use a hybrid ABC-MH algorithm. Specifically, we use an approximate Bayesian computation (ABC) scheme to draw weighted samples of (*c, σ*^2^) in the first stage followed by a Metropolis-Hastings (MH) step that samples (𝒯, ***t***) given ABC samples of (*c, σ*^2^) in the second stage. Motivation for using ABC in the first stage stems from: (i) availability of informative statistics; (ii) generation of better quality samples of the tree (compared to a single-stage MH); and (iii) better computational efficiency. We refer to Section 4.2 for more details.

### 3.1 Hybrid ABC-MH Algorithm

ABC is a family of inference techniques that are designed to estimate the posterior density pr(𝒟 |*θ*) of parameters *θ* given data 𝒟 when the corresponding likelihood pr(𝒟 |*θ*) is intractable but fairly simple to sample from. Summarily, ABC approximates pr(*θ*|*𝒟*) by pr(*θ* |***S***_*obs*_) where ***S***_*obs*_ is a *d*-dimensional summary statistic that ideally captures most information about *θ*. In the special case where ***S***_*obs*_ is a vector of sufficient statistics, it is well known that pr(*θ*|*𝒟*) = pr(*θ* |***S***_*obs*_). To generate a sample from the partial posterior distribution pr(*θ* |***S***_*obs*_), ABC with rejection sampling proceeds by: (i) simulating *N* ^syn^ values *θ*_*l*_, *l* = 1, …, *N* ^syn^ from the prior distribution pr(*θ*); (ii) simulating datasets *D*_*l*_ from pr(𝒟|*θ*_*l*_); (iii) computing summary statistics ***S***_*l*_, *l* = 1, …, *N* ^syn^ from 𝒟_*l*_; (iv) retaining a subset of 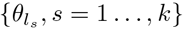 of size *k < N* ^syn^ that corresponds to ‘small’ 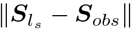 values based on some threshold. Given pairs 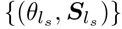, the task of estimating the partial posterior translates to a problem of conditional density estimation, e.g., based on Nadaraya-Waston type estimators and local regression adjustment variants to correct for the fact that 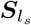 may not be exactly ***S***_*obs*_; see Sisson et al. [2018] for a comprehensive review. To implement ABC, the choice of summary statistics is central.

We detail the specialization of ABC to the marginal posterior distributions of *c* and *σ*^2^ in Section 3.1.1. Given any pair of (*c, σ*^2^), we can sample trees from a density function up to an unknown normalizing constant based on an existing MH algorithm [Knowles and Ghahramani, 2015]. Our proposal is to condition on the posterior median of (*c, σ*^2^) of ABC-weighted samples from the first stage, when sampling the trees in the second stage; clearly, other choices are also available. This strategy produced comparable MAP trees and inference of other tree-derived results relative to tree samples based on full ABC samples of *c* and *σ*^2^.

Pseudo code for the two-stage algorithm is presented in the Supplementary Material Algorithm S1. We briefly describe below its key components.

#### 3.1.1 Stage 1: Sampling Euclidean Parameters (*c, σ*^2^) using ABC

Accuracy and efficiency of the ABC procedure is linked to two competing desiderata on the summary statistics: (i) informative, or ideally sufficient; (ii) low-dimensional.

##### Summary statistic for *σ*^2^

From the closed-form likelihood in Equation (5), a sufficient statistic of *σ*^2^**Σ**^𝒯^ is easily available, using which we construct a summary statistics for *σ*^2^.

###### Lemma 1.

*With* **X** *as the observed data, the statistic* 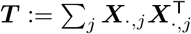*is sufficient for σ*^2^**Σ**^*T*^ *and follows a Wishart distribution W*_*I*_ (*J, σ*^2^**Σ**^𝒯^), *where* ***X***_.,*j*_ = [*x*_1*j*_, …, *x*_*Ij*_] ∈ ℝ ^*I*^. *Then with* 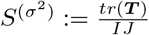 *we have* 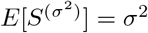 *and* 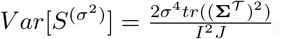

Due to the normality of **X** in (5), and the Factorization theorem [Casella and Berger, 2001], we see that ***T*** is complete and sufficient for *σ*^2^**Σ**^𝒯^ and ***T*** ∼ *W*_*I*_ (*J*, ∼ *σ*^2^**Σ**^𝒯^). Well-known results about the trace and determinant of **X** (see for e.g. Mathai [1980]) provide the stated results on the mean and variance of *tr*(***T***). Owing to its unbiasedness, we choose 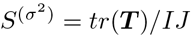 as the summary statistic for *σ*^2^ and examine its performance through simulations in Section 4; other choices are assessed in the Supplementary Material Section S4.1.

##### Summary statistic for *c*

Based on the matrix normal distribution of Proposition 1, the divergence parameter *c* does not appear in the observed data likelihood. Any statistic based on the entire observed data set **X** is sufficient, but not necessarily informative about *c*. In DDT, the prior distribution of the vector of branching times ***t*** is governed by divergence parameter *c* via the divergence function *a*(*t*; *c*). Thus an informative summary statistic for *c* can be chosen by assessing its information about ***t***. For example, tighter observed clusters indicate small *c* (e.g., *c <* 1), where the level of tightness is indicated by the branch lengths from leaves to their respective parents. We construct summary statistics for *c* based on a dendrogram estimated via hierarchical clustering of **X** based on pairwise distances *δ*_*i,i*′_ := ‖ ***X***_*i*_ − ***X***_*i*′_ ‖, *i* =≠*i*′. The summary statistics ***S***^(*c*)^ we choose is a ten-dimensional concatenated vector comprising the 10th, 25th, 50th, 75th and 90th percentiles of empirical distribution of: (i) *δ*_*i,i*′_ ; (ii) branch lengths associated with leaves of the dendrogram. Other candidate summary statistics for *c* are examined in Supplementary Material Section S4.1.

#### 3.1.2 Stage 2: Sampling Tree Parameters (𝒯, *t*) using Metropolis-Hastings

For the second stage, we proceed by choosing a representative value 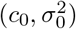 chosen from the posterior sample of (*c, σ*^2^), which in our case is the posterior median. Then a Metropolis-Hastings (MH) algorithm to sample from 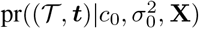; recall that the R_x_tree is characterized by both the topology 𝒯 and divergence times ***t***. In particular, after initialization (e.g., the dendrogram obtained from hierarchical clustering), we first generate a candidate tree (𝒯 ′, ***t***′) from the current tree (𝒯, ***t***) in two steps: (i) detaching a subtree from the original tree; (ii) reattaching the subtree back to the remaining tree. Acceptance probabilities for a candidate tree can be computed exactly and directly using the explicit likelihood in (5), without which they would have to be calculated iteratively [Neal, 2003, Knowles and Ghahramani, 2015]. See Supplementary Material Section S2.2 for details of the proposal function and the acceptance probabilities.

##### Remark 1.

In order to use the explicit likelihood in (5) from Proposition 1 to generate observed data **X**, a tree-structured covariance **Σ**^𝒯^ needs to be specified, whose entries in-turn depend on the parameter *c* through the divergence function.It is not straightforward to fix or sample a **Σ**^𝒯^ since its entries need to satisfy the inequalities (3). It is easier to generate data **X** directly using the DDT generative mechanism in the ABC stage, and this is the approach we follow and is described in Supplementary Section S2.

Summarily, there are three main advantages to using the explicit likelihood from Proposition 1: (i) decoupling of Euclidean and tree parameters to enable an efficient two-stage sampling algorithm; (ii) direct and exact computation of tree acceptance probabilities in MH stage; (iii) determination of informative sufficient statistic for *σ*^2^ (Lemma 1).

### 3.2 Posterior Summary of R_x_-Tree, (𝒯, *t*)

While quantifying uncertainty concerning the tree parameters (𝒯, ***t***) is of main interest, we note that, from definition of the DDT, this is influenced by uncertainty in the model parameters. In particular, the first stage of ABC-MH produces weighted samples and we calculate the posterior median by fitting an intercept-only quantile regression with weights (see details in the Supplementary Material Section S2.1). For the R_x_-tree, we consider global and local tree posterior summaries that capture uncertainty in the latent hierarchy among all and subsets of treatments.

Flexible posterior inference is readily available based on *L* posterior samples of (𝒯, ***t***) from the MH step. It is possible to construct correspond tree-structured covariance matrices **Σ**^𝒯^ from sample (𝒯, ***t***). Instead, we compute:

a. a global *maximum a posteriori* (MAP) estimate of the R_x_-tree that represents the overall hierarchy underlying the treatment responses;
b. local uncertainty estimates of co-clustering probabilities among a subset 𝒜 ⊂ {1, …, *I*} of treatments based on posterior samples of the corresponding subset of divergence times.

#### Posterior co-clustering probability functions

We elaborate on the local summary (b). Suppose 𝒜= {*i, i*′, *i*″} consists of three treatments. Given a tree topology 𝒯, note that at every *t* ∈ [0, 1] a clustering of all *I* treatments is available and the clustering changes only at times 0 *< t*_1_ *< … < t*_*I*−1_. Consequently, for a given tree topology 𝒯drawn from its posterior, we can compute for every level *t* ∈ [0, 1] a posterior probability that *i, i*′ and *i*″ belong to the same cluster. Such a posterior probability can be approximated using Monte Carlo on the *L* posterior samples. Accordingly, we define the estimated posterior co-clustering probability (PCP) function associated with 𝒜 as,

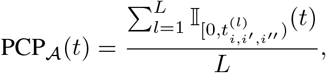

where 𝕀_*B*_ is the indicator function on the set *B* and 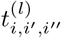 is the divergence time of 𝒜 = {*i, i*′, *i*″} in the *l*-th tree sample. Essentially, the PCP _𝒜_ (*t*) can be viewed as the proportion of tree samples with {*i, i*′, *i*″} having the most recent common ancestor later than *t*.

For every subset 𝒜, the function [0, 1] ∋ *t* ↦ PCP_𝒜_ (*t*) ∈ [0, 1] is non-increasing starting at 1 and ending at 0, and reveals propensity among treatments in 𝒜 to cluster as one traverses down an (estimate of) R_x_-tree starting at the root: a curve that remains flat and drops quickly near 1 indicates higher relative similarity among the treatments in 𝒜 relative to the rest of the treatments. A scalar summary of PCP_𝒜_ (*t*) is the area under its curve known as integrated PCP iPCP_𝒜_, which owing to the definition of PCP_𝒜_ (*t*), can be interpreted as the expected (or average) chance of co-clustering for treatments in 𝒜.

Figure 3 illustrates an example of a three-way iPCP_𝒜_ with 𝒜= {*i, i*′, *i*″} for a PDX data with *I* treatments and *J* patients (Figure 3(A)). Given *L* = 3 posterior trees samples (Figure 3(B)) drawn from the PDX data, we first calculate the whole PCP_𝒜_ (*t*) function by moving the time *t* from 0 to 1. Starting from time *t* = 0, no treatment diverges at time *t* = 0 and the PCP_𝒜_ (*t*) is 1. At time *t*′, treatments diverge in one out of the three posterior trees and PCP_𝒜_ (*t*) therefore drops from 1 to 2*/*3. Moving the time toward *t* = 1, treatments diverge in all trees and the PCP_𝒜_ (*t*) drops to 0. The iPCP_𝒜_ then can be obtained by the area under the PCP_𝒜_ (*t*).

**Figure 3:**
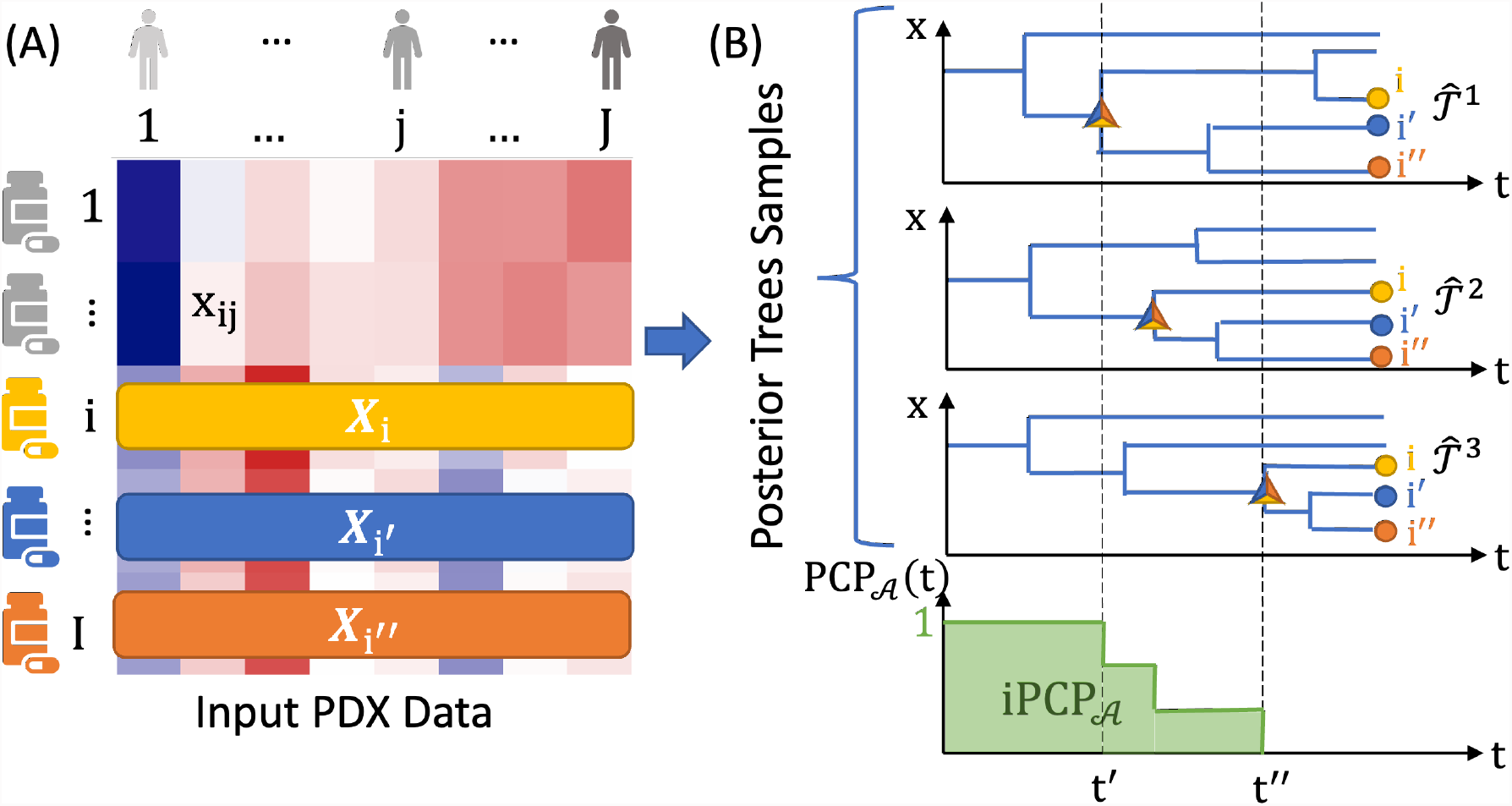
Posterior tree summaries. (A) The input PDX data with *I* treatments and *J* patients, and treatments 𝒜 = {*i, i*′, *i*″} are of interest. (B) PCP_𝒜_ (*t*) and iPCP_𝒜_ for treatments 𝒜 based on *L* = 3 posterior trees. The relevant divergence times are represented by a “△” in each posterior tree sample. For example, at time *t*′, the treatments in 𝒜 diverge in one out of the three trees. Because PCP_𝒜_ (*t*′) is defined by the proportion of posterior tree samples in which 𝒜 has *not* diverged up to and including *t*′, it drops from 1 to 2*/*3.

##### Remark 2.

In the special case of𝒜 = {*i, i*′} for two treatments, the definition of iPCP_𝒜_ can be related to the cophenetic distance [Sokal and Rohlf, 1962, Cardona et al., 2013] and, moreover, extends definition of the cophenetic distance to multiple trees. Given two treatments *i* and *i*′ in a single tree, let *t*_*d*_ be the time at which their corresponding Brownian paths diverge. Then 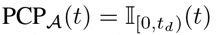 and iPCP_𝒜_ = *t*_*d*_; this implies that the cophenetic distance is 2(1 − *t*_*d*_) and thus iPCP_𝒜_ and the cophenetic distances uniquely determines the same tree structure. For *L >* 1 trees, a Carlo average of divergence times of *L* trees leads to the corresponding iPCP_𝒜_.

##### Remark 3.

Given *I* treatments, since pairwise cophenetic distances from one tree determines a tree [Lapointe and Legendre, 1991, McCullagh, 2006], one might consider summarizing and represent posterior trees in terms of an *I* × *I* matrix **Σ** consisting of entries iPCP{_*i,i*′_*}* for every pair of treatments of (*i, i*′), estimated from the posterior sample of trees. However, **Σ** need not to be a tree-structured matrix that uniquely encodes a tree. It is possible to project **Σ** on to the space of tree-structured matrices (see for e.g., Bravo et al. [2009]) but the projection might result in a non-binary tree structure. We discuss this issue and its resolution in Supplementary Material Section S3.

## 4 Simulations

Accurate characterization of similarities among any subset of treatments is central to our scientific interest in identifying the promising treatment subsets for further investigation. In addition, we have introduced a two-stage algorithm to improve our ability to efficiently draw tree samples from the posterior distribution (similarly for the Euclidean parameters). To demonstrate the modeling and computational advantages, we conduct two sets of simulations. The first simulation shows that the proposed model estimates the similarity (via iPCP) better than alternatives, even when the true data generating mechanisms deviate from DDT assumptions in terms of the form of divergence function, prior distribution for the unknown tree, and normality of the responses. The second simulation illustrates the computational efficiency of the proposed two-stage algorithm in producing higher quality posterior samples of Euclidean parameters, resulting in more accurate subsequent estimation of an unknown tree and iPCPs, two key quantities to our interpretation of real data results.

### 4.1 Simulation I: Estimating Treatment Similarities

We first show that iPCPs estimated by DDT are closer to the true similarities (operationalized by functions of elements in the true divergence times in **Σ**^𝒯^) under different true data generating mechanisms that may follow or deviate from the DDT model assumptions in three distinct aspects (the form of divergence function, the prior distribution over the unknown tree, and normality).

#### Simulation setup

We simulate data by mimicking the PDX breast cancer data (see Section 5) with *I* = 20 treatments and *J* = 38 patients. We set the true scale parameter as the posterior median 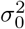 and the true tree 𝒯_0_ as the MAP tree that are estimated from the breast cancer data; We consider four scenarios to represent different levels of deviation from the DDT model assumptions:

i. No deviation of the true data generating mechanism from the fitted DDT models: given 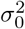 and 𝒯_0_, simulate data based on the DDT marginal data distribution (Equation (5));
The true data generating mechanism deviates from the fitted DDT in terms of:
ii. divergence function: same as in (i), but the true tree is a random tree from DDT with misspecified divergence function, 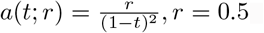;
iii. prior for tree topology: same as in (i), but the true tree is a random tree from the coalescence model (generated by function `rcoal` in R package `ape`), and,
iv. marginal data distribution: same as in (i), but the marginal likelihood is a centered multivariate *t* distribution with degree-of-freedom four and scaled by 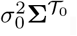.

For each of four true data generating mechanisms above, we simulate *B* = 50 replicate data sets. In the following, we use the DDT model and the two-stage algorithm for all estimation regardless of the true data generating mechanisms. For DDT, we ran the two-stage algorithm where the second stage is implemented with five parallel chains. For each chain, we ran 10, 000 iterations, discarded first 9, 000 trees and combined five chains with a total of 5, 000 posterior tree samples.

First, we compute the iPCPs for all pairs of treatment combinations following the definition of iPCP_𝒜_ where 𝒜={ *i, i*′ }, 1≤ *i < i*′ ≤*I*. Two alternative approaches to defining and estimating similarities between treatments are considered: (i) similarity derived from agglomerative hierarchical clustering, and (ii) empirical Pearson correlation of the two vectors of responses ***X***_*i*_ and ***X***_*i*′_, for *i* ≠*i*^*′*^ In particular, for (i), we considered five different linkage methods (Ward, Ward’s D2, single, complete and Mcquitty) with Euclidean distances. Given an estimated dendrogram from hierarchical clustering, the similarity for a pair of treatments is defined by first normalizing the sum of branch lengths from the root to leaf as 1, and then calculating the area under of the co-clustering curve (AUC) obtained by cutting the dendrogram at various levels from 0 to 1. For three- or higher-way comparisons, (i) can still produce an AUC based on a dendrogram obtained from hierarchical clustering, while the empirical Pearson correlation in (ii) is undefined hence not viable as a comparator beyond assessing pairwise treatment similarities.

#### Performance metrics

For treatment pairs 𝒜 = {*i, i*′ *}*, to assess the quality of estimated treatment similarities for each of the methods above (DDT-based, hierarchical-clustering-based, and empirical Pearson correlation), we compare the estimated values against the true branching time 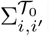; similarly when assessing recovery of three-way treatment similarities, e.g., 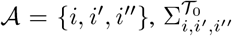 is defined as the time when {*i, i*′, *i*″} first branches in the true tree 𝒯_0_. In particular, for replication data set *b* = 1, …, *B*, let 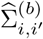 generically represent the pairwise similarities for treatment subsets (*i, i*′) that can be based on DDT, hierarchical clustering or empirical pairwise Pearson correlation. For three-way comparisons, let 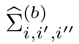 generically represent the three-way similarities for treatment subset (*i, i*′, *i*″) that can be based on DDT, or hierarchical clustering.

We assess the goodness of recovery by computing 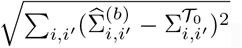, the Frobenious norm of the matrix in recovering the entire 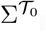. We compute 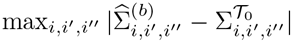, the max-norm of the matrix in recovering the true three-way similarities. For a given method and treatment subset 𝒜, the above procedure results in *B* values, the distribution of which can be compared across methods; smaller values indicate better recovery of the true similarities.

Alternatively, for each method and each treatment subset, we also compute the Pearson correlation between the estimated similarities and the true branching times across replicates for pairwise or three-way treatment subsets: 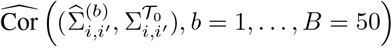, for treatments *i < i*′ and 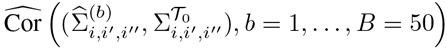, for treatments *i < i*′ *< i*″. We refer to this metric as “Correlation of correlations” (the latter uses the fact that the entries in the true 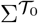 being correlations; see Equation (5)); higher values indicate better recovery of the true similarities.

#### Simulation results

We observe that DDT better estimates the treatment similarities even under misspecified models. In particular, under scenarios where the true data generating mechanisms deviate from the fitted DDT model assumptions (ii-iv), the DDT captures the true pairwise and three-way treatment similarities the best by higher values in correlation of correlations (left panels, Figure 4) and lower matrix/array distances (right panels, Figure 4). In particular, the fitted DDT with divergence function *a*(*t*) = *c/*(1− *t*) under Scenario i, ii and iii performed similarly well indicating the relative insensitivity to the DDT modeling assumptions with respect to divergence function and the tree generative model. Under Scenario iv where the marginal likelihood assumption deviates from Gaussian with heavier tails, the similarity estimates from all methods deteriorate relative to Scenarios i-iii. Comparing between methods, the similarities derived from hierarchical clustering with single linkage is comparable to DDT model when evaluated by correlation of correlation, but worse than DDT when evaluated by the matrix norm.

**Figure 4:**
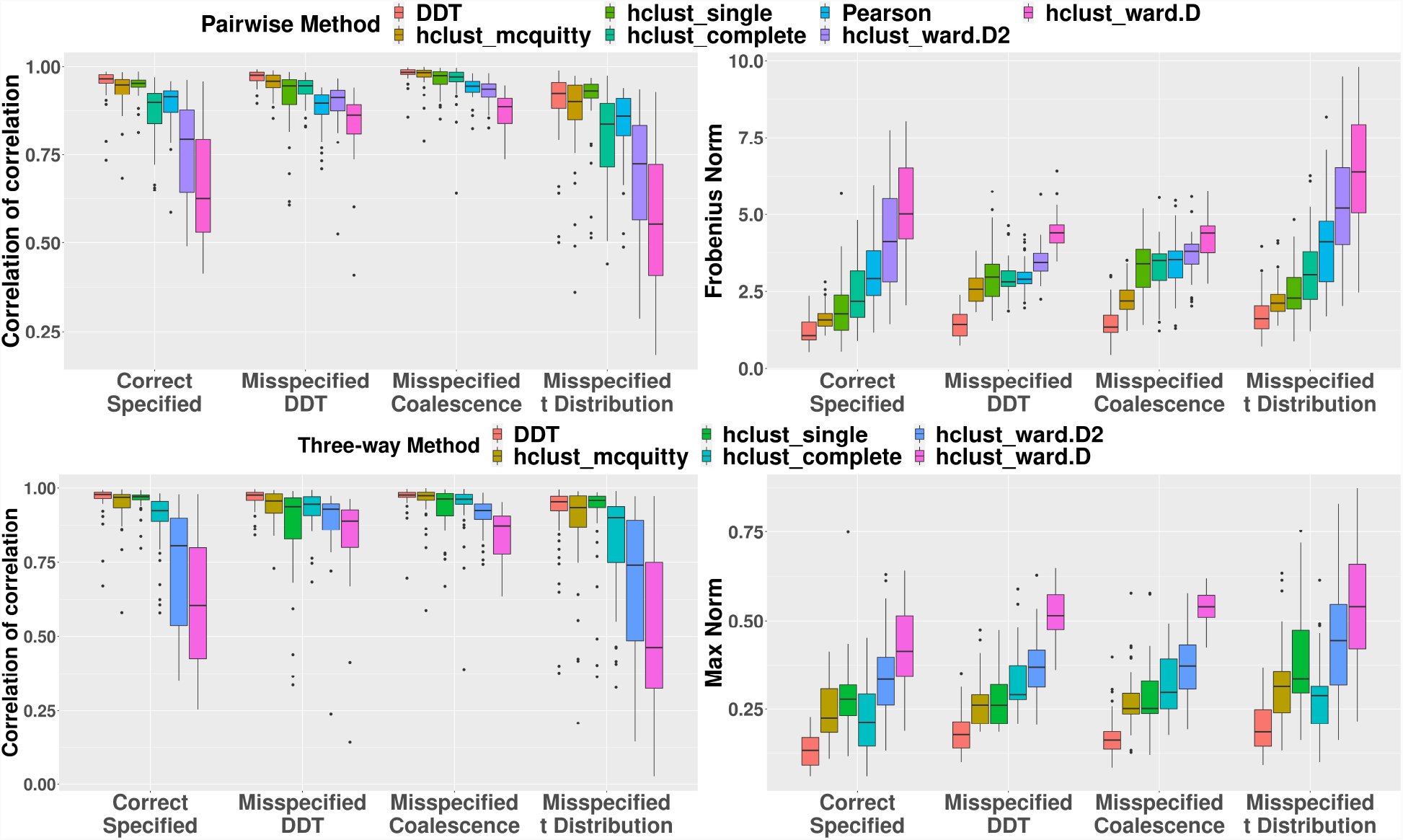
Simulation studies for comparing the quality of estimated treatment similarities based on DDT, hierarchical clustering, and empirical Pearson correlation. Two performance metrics are used: (Left) Correlation of correlation (higher values are better); (Right) Matrix distances with Frobenius norm for pairwise similarity and max norm for three-way similarity (lower values are better). DDT captures both true pairwise (upper panels) and three-way (lower panels) similarity best under four levels of misspecification scenarios.

### 4.2 Simulation II: Comparison with Single-Stage MCMC Algorithms

We have also conducted extensive simulation studies that focus on the computational aspect of the proposed algorithms and demonstrate the advantage of the proposed two-stage algorithm in producing higher quality posterior samples of the unknown tree than classical single-stage MCMC algorithms. In particular, we demonstrate that the proposed algorithm produces (i) MAP trees that are closer to the true tree than alternatives (hierarchical clustering, single-stage MH with default hierarchical clustering or the true tree at initialization) and (ii) more accurate estimation of pairwise treatment similarities compared to single-stage MCMC algorithms. See Supplementary Material Section S5 for further details.

#### Additional simulations and sensitivity analyses

Aside from the simulations above focusing on the tree structure and the divergence time, Supplementary Material S4 offers additional details for Euclidean parameters including the parameter inference and algorithm diagnostics. In particular, we empirically show that current ***S***^(*c*)^ and 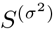 outperform other candidate summary statistics in terms of bias in Section S4.1. In Section S4.2, we present additional simulation results that demonstrate that the two-stage algorithm (i) enjoys stable effective sample size (ESS) for (*c, σ*^2^); (ii) leads to similar or better inference on (*c, σ*^2^), as ascertained using credible intervals. Section S4.3 checks the convergence of MH and the goodness of fit for ABC.

## 5 Treatment Trees in Cancer using PDX Data

### 5.1 Dataset Overview and Key Scientific Questions

We leverage a recently collated PDX dataset from the <u>N</u>ovartis <u>I</u>nstitutes for <u>B</u>ioMedical <u>R</u>esearch - <u>PDX E</u>ncyclopedia [NIBR-PDXE, [Gao et al., 2015]] that interrogated multiple targeted therapies across different cancers and established that PDX systems provide a more accurate measure of the response of a population of patients than traditional preclinical models. Briefly, the NIBR-PDXE consists of *>* 1, 000 PDX lines across a range of human cancers and uses a 1× 1 × 1 design (one animal per PDX model per treatment); i.e., each PDX line from a given patient was treated simultaneously with multiple treatments allowing for direct assessments of treatment hierarchies and responses. In this paper, we focus on our analyses on a subset of PDX lines with complete responses across five common human cancers: Breast cancer (BRCA), Cutaneous Melanoma (CM, skin cancer), Colorectal cancer (CRC), Non-small Cell Lung Carcinoma (NSCLC), and Pancreatic Ductal Adenocarcinoma (PDAC). After re-scaling data and missing data imputation, different numbers of treatments, *I*, and PDX models, *J*, presented in the five cancers were, (*I, J*): BRCA, (20, 38); CRC, (20, 40); CM, (14, 32); NSCLC, (21, 25); and PDAC, (20, 36). (See Supplementary Material Table S7 for treatment names and Section S6.1 for details of pre-processing procedures.)

In our analysis, we used the best average response (BAR) as the main response, by taking the untreated group as the reference group and using the tumor size difference before and after administration of the treatment(s) following Rashid et al. [2020]. Positive values of BAR indicate the treatment(s) shrunk the tumor more than the untreated group with higher values indicative of (higher) treatment efficacy. The treatments included both drugs administered individually with established mechanisms (referred to as “monotherapy”) and multiple drugs combined with potentially unknown synergistic effects (referred to as “combination therapy”). Our key scientific questions were as follows: (a) identify plausible biological mechanisms that characterize treatment responses for monotherapies within and between cancers; (b) evaluate the effectiveness of combination therapies based on biological mechanisms. Due to a potentially better outcome and lower resistance, combination therapy with synergistic mechanism is highly desirable [Bayat Mokhtari et al., 2017].

#### DDT model setup

For all analyses we followed the setup in the Section 4.1 and obtained *N* ^syn^ = 600, 000 synthetic datasets from the ABC algorithm (Section 3.1.1) with prior *c* ∼ Gamma(2, 2) and 1*/σ*^2^ ∼Gamma(1, 1) and took the first 0.5% (*d* = 0.5%) closest data in terms of ***S***^(*c*)^ and 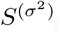 We calculated the posterior median of (*c, σ*^2^) as described in Section 3.2. For the second-stage MH, we ran five chains of the two-stage algorithm with (*c, σ*^2^) fixed at the posterior median by 10,000 iterations and discarded the first 9,000 trees, which resulted in 5,000 posterior trees in total. Finally, we calculated the R_x_-tree (MAP) and iPCP based on 5,000 posterior trees for all subsequent analyses and interpretations. All computations were divided on multiple different CPUs (see the Supplementary Table S5 for the full list of CPUs). For the BRCA data with *I* = 20 and *J* = 38, we divided the ABC stage into 34 compute cores with a total of 141 CPU hours and maximum 4.7 hours in real time. For the MH stage and the single-stage MCMC, we split the computation on 5 compute cores with a total of 8.6 and 12 CPU hours, and a maximum 1.7 and 2.5 hours in real time, respectively.

Our results are organized as follows: we provide a summary of the R_x_-tree estimation and treatment clusters in Section 5.2 followed by specific biological and translational interpretations in Sections 5.3 and 5.4 for monotherapy and combination therapy, respectively. Additional results can be accessed and visualized using our companion R-shiny application (see Supplementary Material Section S6.4 for details).

### 5.2 R_x_-tree Estimation and Treatment Clusters

We focus our discussion on three cancers: BRCA, CRC and CM here – see Supplementary Materials Section S6.3 for NSCLC and PDAC. In Figure 5, R_x_-tree, pairwise iPCP and (scaled) Pearson correlation are shown in the left, middle and right panels, respectively. Focusing on the left two panels, we observe that the R_x_-tree and the pairwise iPCP matrix show the similar clustering patterns. For example, three combination therapies in CM form a tight subtree and are labeled by a box in the R_x_-tree of Figure 5 and a block with higher values of iPCP among three combination therapies also shows up in the corresponding iPCP matrix with a box labeled. In our analysis, the treatments predominantly target six oncogenic pathways that are closely related to the cell proliferation and cell cycle: (i) phosphoinositide 3-kinases, PI3K; (ii) mitogen-activated protein kinases, MAPK; (iii) cyclin-dependent kinases, CDK; (iv) murine double minute 2, MDM2; (v) janus kinase, JAK; (vi) serine/threonine-protein kinase B-Raf, BRAF. We label targeting pathways above for monotherapies with solid dots and further group PI3K, MAPK and CDK due to the common downstream mechanisms [e.g., Repetto et al., 2018, Kurtzeborn et al., 2019]. Roughly, the R_x_-tree from our model clusters monotherapies targeting oncogenic processes above and largely agrees with common and established biology mechanisms. For example, all PI3K-MAPK-CDK inhibitors (solid square) belong to a tighter subtree in three cancers; two MDM2 monotherapies (solid triangle) are closest in both BRCA and CRC. While visual inspection of the MAP R_x_-tree agrees with known biology, iPCP further quantifies the similarity by assimilating the information across multiple trees from our MCMC samples. For the ensuing interpretations in Sections 5.3 and 5.4, we focus on iPCP and verify our model through monotherapies with known biology, since our a priori hypothesis is that monotherapies that share the same downstream pathways should exhibit higher iPCP values. Furthermore, we extend our work to identify combination therapies with synergy and discover several combination therapies for each cancer.

**Figure 5:**
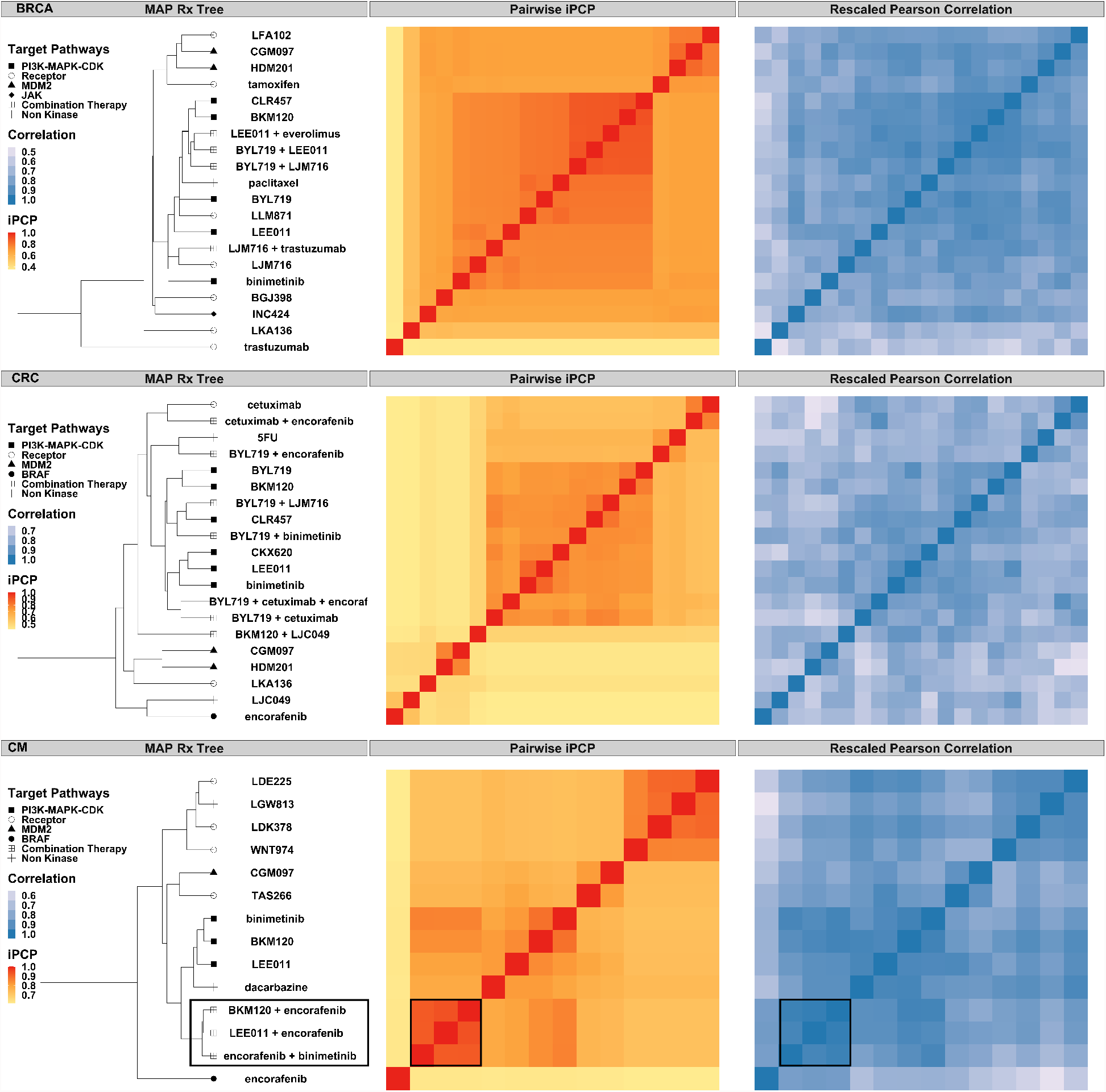
The R_x_-tree and iPCP for breast cancer (BRCA, top row), colorectal cancer (CRC, middle row) and melanoma (CM, lower row). Three panels in each row represent: (left) estimated R_x_-tree (MAP); distinct external target pathway information is shown in distinct shapes for groups of treatments on the leaves; (middle) estimated pairwise iPCP, i.e., the posterior mean divergence time for pairs of entities on the leaves (see the result paragraph for definition for any subset of entities); (right) scaled Pearson correlation for each pair of treatments. Note that the MAP visualizes the hierarchy among treatments; the iPCP is not calculated based on the MAP, but based on posterior tree samples (see definition in Section 3.2)

### 5.3 Biological Mechanisms in Monotherapy

Our estimation procedure exhibits a high level of concordance between known biological mechanisms and established monotherapies for multiple key signalling pathways. From the R_x_-tree in Figure 5, aside from the oncogenic process (solid dots) introduced above, monotherapies also target receptors (hollow circles) or other non-kinase targets (e.g. tubulin; crosses). We summarize our key findings and interpretations for each of signaling pathways and their regulatory axes, namely, PI3K-MAPK-CDK and MDM2 from cell cycle regulatory pathways, human epidermal growth factor receptor 3 (ERBB3) from receptor pathways, and tubulin from non-kinase pathways along with their implications in monotherapy across different cancers.

#### PI3K-MAPK-CDK inhibitors

For treatments targeting PI3K, MAPK and CDK, treatments have the same target share high iPCP. In the NIBR-PDXE dataset, three PI3K inhibitors (BKM120, BYL719 and CLR457), two MAPK inhibitors (binimetinib and CKX620) and one CDK inhibitor (LEE011) were tested, but different cancers contain different numbers of treatments. Specifically, all three PI3K inhibitors present in BRCA and CRC, but only BKM120 is tested in CM; CRC contains two MAPK inhibitors while BRCA and CM only have binimetinib; LEE011 is tested in all three cancers. In Figure 6, BKM120, BYL719 and CLR457 share high pairwise iPCPs (box (1)) and all target PI3K for BRCA and CRC (BRCA, (BKM120, CLR457): 0.8986, (BKM120, BYL719): 0.8002, (BYL719, CLR457): 0.8002; CRC, (BKM120, CLR457): 0.7555, (BKM120, BYL719): 0.8041, (BYL719, CLR457): 0.7597); MAPK (box (2)) inhibitors, binimetinib and CKX620, show a high pairwise iPCP in CRC (0.7792). Asides from the pairwise iPCPs, our model also suggests high multi-way iPCPs among PI3K inhibitors in BRCA (0.8002) and CRC (0.7513). Among these inhibitors, PI3K inhibitor of BYL719 was approved by FDA for breast cancer; MAPK inhibitor of binimetinib was approved by FDA for BRAF mutant melanoma in combination with encorafenib; and CDK inhibitor of LEE011 was approved for breast cancer.

**Figure 6:**
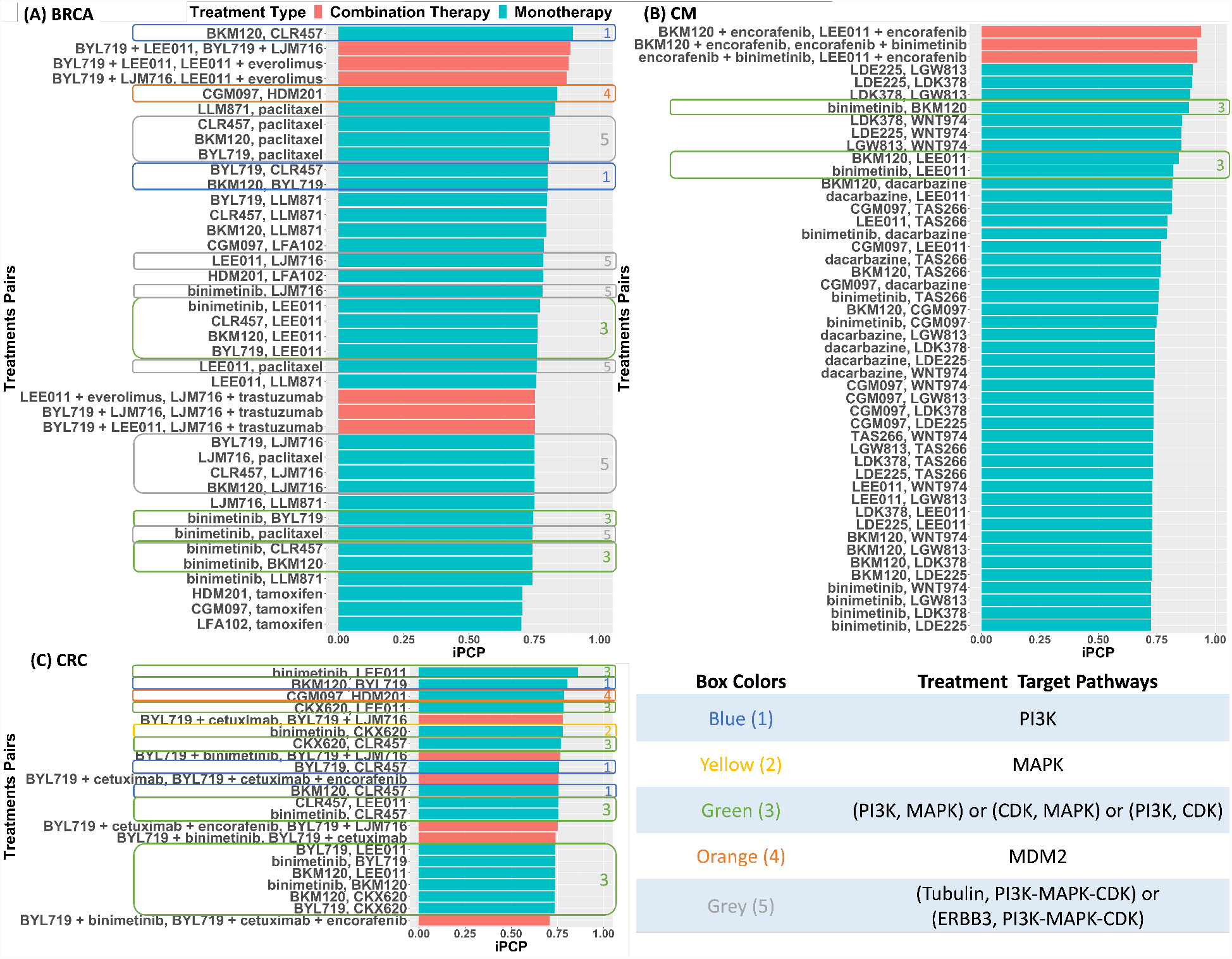
Bar plot of iPCPs for pairs of combination therapies (red bars) and pairs of monotherapies (green bars): (A) breast cancer, (B) melanoma and (C) colorectal cancer. The bar plots are sorted by the iPCP values (high to low); pairs of treatments are shown only if the estimated iPCP is greater than 0.7. Monotherapies have different known targets which are listed in the bottom-right table (see Section 5.3 for more details and discussion on monotherapies).

Our model suggests treatments targeting different pathways also share high iPCP values across different cancers. Monotherapies targeting different cell cycle regulatory pathways (PI3K, MAPK and CDK) exhibit high iPCPs. CDK inhibitor, LEE011, and MAPK inhibitors share high pairwise iPCP values in BRCA ((LEE011, binimetinib): 0.7709), CRC ((LEE011, binimetinib): 0.8617, (LEE011, CKX620): 0.7820) and CM ((LEE011, binimetinib): 0.8210) in the Figure 6 with box (3). High iPCP among MAPK and CDK inhibitors agree with biology, since it is known that CDK and MAPK collaboratively regulate downstream pathways such as Ste5 [Repetto et al., 2018]. High pairwise iPCP values between PI3K and MAPK inhibitors were observed in box (3) in the Figure 6. Specifically, our model suggests high pairwise iPCPs as follows: (i) BRCA, (binimetinib, BKM120): 0.7427, (binimetinib, BYL719): 0.7441, (binimetinib, CLR457): 0.7427)); (ii) CRC, (binimetinib, BKM120): 0.7374, (binimetinib, BYL719): 0.7388, (binimetinib, CLR457): 0.7541, (CKX620, BKM120): 0.7366, (CKX620, BYL719): 0.7357, (CKX620, CLR457): 0.7676)); (iii) CM, (binimetinib, BKM120): 0.8882. Aside from the pairwise iPCPs above, high multi-way iPCPs in BRCA (0.7422), CRC (0.7300) and CM (0.8882) also show the similar information. From the existing literature, both PI3K and MAPK can be induced by ERBB3 phosphorylation [Balko et al., 2012] and it is not surprising to see high iPCPs between PI3K and MAPK inhibitors.

#### ERBB3 and tubulin inhibitors

Our model also found high iPCP values among ERBB3, tubulin and PI3K-MAPK-CDK inhibitors in BRCA. ERRB3 inhibitor, LJM716, exhibits high pairwise iPCP values with PI3K (BKM120: 0.7501, BYL719: 0.7513, CLR457: 0.7500), MAPK (binimetinib: 0.7811), CDK (LEE011: 0.7847) and tubulin (paclitaxel: 0.7505) inhibitors in the Figure 6 with box (5). Since PI3K and MAPK are downstream pathways of ERBB3 [Balko et al., 2012] and CDK works closely with PI3K and MAPK [Kurtzeborn et al., 2019, Repetto et al., 2018], high iPCPs between ERBB3 inhibitor and PI3K-MAPK-CDK inhibitors are not surprising. For ERBB3 and tubulin, ERBB3 is a critical regulator of microtubule assembly [Wu et al., 2021] and tubulin plays an important role in building microtubules. Since microtubules form the skeletons of cells and are essential for cell division [Gunning et al., 2015, Haider et al., 2019], tubulin inhibitor, paclitaxel, kills cancer cell by interfering cell division and is an FDA-approved treatment. In congruence with the above results, tubulin inhibitor paclitaxel also shares high iPCPs with PI3K (BKM120: 0.8076, BYL719: 0.8063, CLR457: 0.8076), MAPK (binimetinib: 0.7433), CDK (LEE011: 0.7587) and ERBB3 (LJM716: 0.7505) in the Figure 6 with box (5). In addition, another CDK4 inhibitor BPT also inhibits tubulin [Mahale et al., 2015] and PI3K inhibitor BKM120 inhibits the formation of microtubule [Bohnacker et al., 2017]. Both offer additional reasons for high iPCP between tubulin and PI3K-MAPK-CDK inhibitors.

#### MDM2 inhibitors

We found two drugs: CGM097 and HDM201 share high iPCP values in BRCA (0.8365) and CRC (0.7860) in the Figure 6 with box (4). Since CGM097 and HDM201 target the same pathway, MDM2, high iPCPs suggest a high similarity between CGM097 and HDM201 and show consistent results between our model and underlying biological mechanism. MDM2 negatively regulates the tumor suppressor, p53 [Zhao et al., 2014] and if MDM2 is suppressed by inhibitors, p53 is able to prevent tumor formation. Both CGM097 and HDM201 entered phase I clinical trial [Konopleva et al., 2020] for wild-type p53 solid tumors and leukemia, respectively.

### 5.4 Implications in Combination Therapy

Based on the concordance between the monotherapy and biology mechanism, we further investigate combination therapies to identify mechanisms with synergistic effect. In NIBR-PDXE, 21 combination therapies were tested and only one of them includes three monotherapies (BYL719 + cetuximab + encorafenib in CRC) and the rest contain two monotherapies. Out of 21 combination therapies, only three do not target any cell cycle (PI3K, MAPK, CDK, MDM2, JAK and BRAF) pathways (see Supplementary Material Table S8 for the full list of combination therapies). From the R_x_-tree in Figure 5, combination therapies tend to form a tighter subtree and are closer to monotherapies targeting PI3K-MAPK-CDK, which implies that the mechanisms under combination therapies are similar to each other and are closer to the PI3K-MAPK-CDK pathways. We identified several combination therapies with known synergistic effects and provide a brief description for each of the cancers in the following paragraphs.

#### Breast cancer

Four combination therapies were tested in BRCA and three therapies targeting PI3K-MAPK-CDK (BYL719 + LJM716, BYL719 + LEE011 and LEE011 + everolimus) form a subtree in R_x_-tree with a high three-way iPCP (0.8719). Among these combination therapies, PI3K-CDK inhibitor, BYL719 + LEE011, suggests a possible synergistic regulation [Vora et al., 2014, Bonelli et al., 2017, Yuan et al., 2019]. Based on the high iPCP between BYL719 + LEE011 and the rest two therapies, we suggest synergistic effect for combination therapies targeting PI3K-ERBB3 (BYL719 + LJM716), and CDK-MTOR (LEE011 + everolimus) for future investigation.

#### Colorectal cancer

Our model suggests a high three-way iPCP (0.7437) among PI3K-EGFR (BYL719 + cetuximab), PI3K-EGFR-BRAF (BYL719 + cetuximab + encorafenib) and PI3K-ERBB3 (BYL719 + LJM716) inhibitors. Since the triple therapy (BYL719 + cetuximab + encorafenib) enters the phase I clinical trial with synergy [Geel et al., 2014], our model proposes the potential synergistic effect for PI3K-ERBB3 based on iPCP for future investigation. Of note, we found a modest iPCP (0.6280) between the FDA-approved combination therapy EGFR-BRAF (cetuximab + encorafenib) and PI3K-EGFR-BRAF (BYL719 + cetuximab + encorafenib) and the modest iPCP can be explained by an additional drug-drug interaction between BYL719 and encorafenib in triple-combined therapy [van Geel et al., 2017].

#### Melanoma

In NIBR-PDXE, three combination therapies were tested in CM, and all of them consist one monotherapy targeting PI3K-MAPK-CDK and the other one targeting BRAF. A tight subtree is observed in the R_x_-tree and our model also suggests a high iPCP (0.9222) among three combination therapies. Since PI3K, MAPK and CDK work closely and share a high iPCP (0.8204) among monotherapies in CM, a high iPCP (0.9222) among three combination therapies is not surprising. Since two combination therapies of BRAF-MAPK (dabrafenib + trametinib and encorafenib + binimetinib) are approved by FDA for BRAF-mutant metastatic melanoma [Dummer et al., 2018a,b, Robert et al., 2019], we recommend the synergy for BRAF-PI3K (encorafenib + BKM120) and BRAF-CDK (encorafenib + LEE011) inhibitors.

### Comparison to alternative approaches

Unlike the probabilistic generative modeling approach proposed in this paper, standard distance-based agglomerative hierarchical clustering and Pearson correlation can also be applied to the PDX data to estimate the similarity. However, simple pairwise similarities can be potentially noisy and the uncertainty in the estimation is not fully incorporated due to the absence of a generative model. As we showed in the Section 4.1 (Simulation I) that agglomerative hierarchical clustering and the Pearson correlation leads to inferior recovery of the true branching times and the true tree structure under different data generating mechanisms mimicking the real data. As further evidence, we compute pairwise similarities based on Pearson correlation (other distance metrics show similar patterns) in the right panel of Figure 5. By mapping the original Pearson correlation *ρ* ∈ [− 1, 1] through a linear function 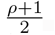, we make the range of iPCP and Pearson correlation comparable. We observe that pairwise iPCP estimated through the DDT model is less noisy than Pearson correlation. For example, both iPCP and Pearson correlation in CM show higher similarities among combination therapy framed by a box, but iPCP exhibits a clearer pattern than Pearson correlation.

## 6 Summary and Discussion

In translational oncology research, PDX studies have emerged as a unique study design that evaluates multiple treatments when applied to samples from the same human tumor implanted into genetically identical mice. PDX systems are promising tools for large-scale screening to evaluate a large number of FDA-approved and novel cancer therapies. However, there remain scientific questions concerning how distinct treatments may be synergistic in inducing similar efficacious responses, and how to identify promising subsets of treatments for further clinical evaluation. To this end, in this paper, we propose a probabilistic framework to learn treatment trees (R_x_-trees) from PDX data to identify promising treatment combinations and plausible biological mechanisms that confer synergistic effect(s). In particular, in a Bayesian framework based on the Dirichlet Diffusion Tree, we estimate a *maximum a posteriori* rooted binary tree with the treatments on the leaves and propose a posterior uncertainty-aware similarity measure (iPCP) for any subset of treatments. The divergence times of the DDT encode the tree topology and are profitably interpreted within the context of an underlying plausible biological mechanism of treatment actions.

From the class of probabilistic models with an unknown tree structure component, we have chosen the DDT mainly owing to the availability of a closed-form marginal likelihood that directly links the tree topological structure to the covariance structure of the observed PDX data, which additionally decouples the Euclidean and tree parameters; to the best of our knowledge this method has not been proposed or explored hitherto for the DDT. The decoupling leads to efficient posterior inference via a two-stage algorithm that confers several advantages. The algorithm generates posterior samples of Euclidean parameters through approximate Bayesian computation and passes the posterior medians to a second stage classical Metropolis-Hastings algorithm for sampling from the conditional posterior distribution of the tree given all other quantities. Through simulation studies, we show that the proposed two-stage algorithm generates better posterior tree samples and captures the true similarity among treatments better than alternatives such as single-stage MCMC and naive Pearson correlations. The posterior samples of trees are summarized by iPCP, which we propose to measure the empirical mechanistic similarity for multiple treatments incorporating uncertainty.

Using the proposed methodology on NIBR-PDXE data, we estimate R_x_-trees and iPCPs for five cancers. Among the monotherapies, iPCP is highly concordant with known biology across different cancers. For example, CGM097 and HDM201 show a high iPCP value among treatments in breast cancer and melanoma, which corroborates known mechanisms, since both monotherapies target the same biological pathway, MDM2, and have been further validated in recent clinical trials [Konopleva et al., 2020]. The proposed iPCP can also suggest improvements upon an existing combination therapy. We first identify a combination therapy with known synergy (not based on the our data) and then determine which additional therapies (monotherapies or combination therapies) have high iPCPs when considered together with the existing combination therapy. Based on the NIBR-PDXE data, for each cancer, we suggest potential synergies between PI3K-ERBB3 and CDK-MTOR for breast cancer, PI3K-ERBB3 for colorectal cancer, and BRAF-PI3K and BRAF-CDK for melanoma that could be potentially explored in future translational studies.

The present analysis shows the promise in guiding the potential treatment strategies by building trees upon NIBR-PDXE dataset using treatment responses, but assume independent patients without using the underlying patient-specific genomic information that is available in the NIBR-PDXE. By including patient-specific genomic information, we may further improve our ability to identify synergistic treatments that may be specific to a subset of patients. One approach to utilize genomic information could be to extend the DDT model to incorporate patient-specific genomic information in the mean structure or the column covariance of the marginal likelihood of Equation (4). Models with non-Gaussian marginal likelihood and non-binary treatment tree in principle can be defined by considering generative tree models based on general diffusion processes [Heaukulani et al., 2014, Knowles and Ghahramani, 2015]. Both extensions raise significant, non-trivial methodological and computation issues (e.g., deriving tractable likelihoods; finding low-dimensional summary statistics for new parameters) and constitute the foundation for future work.

## Code and data availability

We also provide a general purpose code in R that accompanies this manuscript along with all the necessary documentation and datasets required to replicate our results (see https://github.com/bayesrx/RxTree). Furthermore, to aid access and visualization of the results, we have also developed an R-shiny application (see Supplementary Material Section S6.4).

## S1 Proof of Proposition 1

We provide a proof for a tree with four leaves (see Figure S1) and extension to trees with a larger number of leaves follows by induction. The main idea is to merge subtrees backward and integrate out responses of internal nodes when merging subtrees.

*Proof*. Consider a subtree 𝒯 rooted at 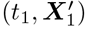 with two leaves (1, ***X***_1_) and (1, ***X***_2_), and one internal node 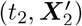 (see Panel (A) of Figure S1). Assume that the root 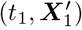 of the subtree is fixed, and responses 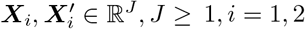, *i* = 1, 2. With ***t*** = (*t*_1_, *t*_2_, *t*_3_)^T^, the conditional distribution for leaf responses would be 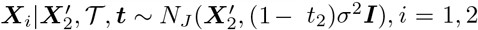. Since 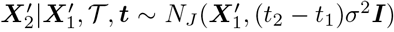, based on the conjugacy of the normal distribution, the marginal distribution is also normal. Conditional on ***t*** and 𝒯, mean and covariance of ***X***_*i*_, *i* = 1, 2 can be derived by the law of iterated expectations and results in the distribution of the subtree 𝒯 ′ with two leaves:

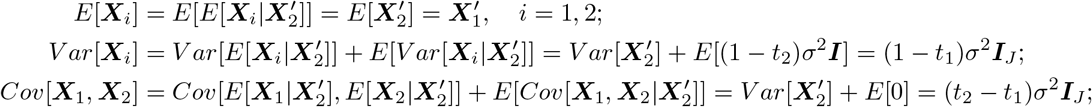

The marginal distribution for the subtree 𝒯 ′ with two leaves is

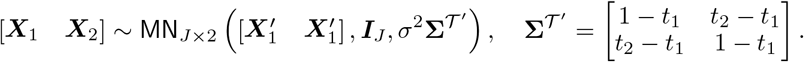

Therefore, we can merge two leaves responses ***X***_1_ and ***X***_2_. Similarly, we can also merge the other subtree 𝒯 ″ to obtain.

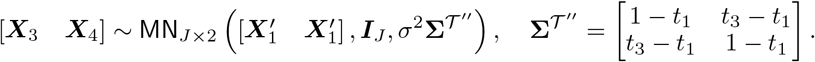

Eventually, we can merge two subtrees (see Panel (B) of Figure S1), 𝒯 ′ and 𝒯 ″. From conjugacy of the normal distribution, the resulting joint marginal distribution of ***X***_*i*_, *i* = 1, 2, 3, 4 is normal. The mean and the variance can be derived along identical lines as above. The only term left is the covariance, and we need to (re-)compute them for locations within and between the combined subtrees. Explicitly,

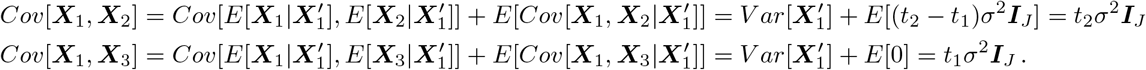

This ensures that

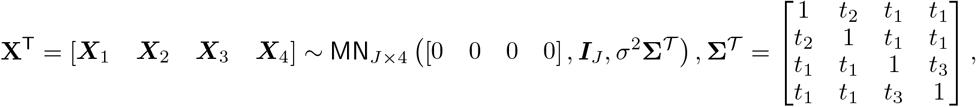

as required. Moreover, denote *t*_*i,i′*_ as the most recent divergence time of leaves *i* and *i′*. We observe that *t*_1_ = *t*_1,3_ = *t*_1,4_ = *t*_2,3_ = *t*_2,4_, *t*_2_ = *t*_1,2_, and *t*_3_ = *t*_3,4_ and complete the Proposition 1. □

**Figure S1:**
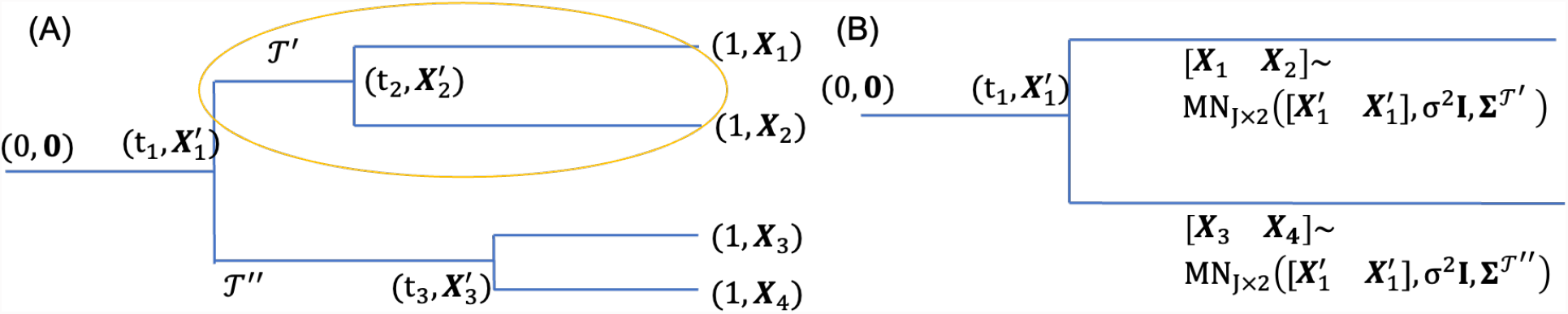
Merging subtrees for the integration process. (A) First step of merging upper subtree, and (B) Final step of merging all subtrees.

## S2 Efficient Two-Stage Hybrid ABC-MH Algorithm

Here we offer details of two-stage algorithm with pseudo code. In the Section S2.1, we describe the full algorithm of the ABC with the following posterior summary of Euclidean parameters (*c, σ*^2^). The Section S2.2 includes the implementation of the proposal function and the acceptance probability of MH stage. Pseudo code for the full two-stage algorithm is presented below in Algorithm S1

### S2.1 ABC Stage and the Posterior Summary of *c* and *σ*^2^

The Section 3 of the Main Paper states the main idea of ABC and we offer the full algorithm of ABC including (i) the synthetic data generation process, (ii) the regression adjustment [Blum, 2010] of ABC, and (iii) posterior summary of the Euclidean parameters.

#### Data generation in ABC

Following Section 2 in the Main Paper, a synthetic data is generated from DDT as follows: (i) given *c*_*l*_ ∼ Gamma(*a*_*c*_, *b*_*c*_), generate a tree 𝒯_*l*_ through the divergence function *a*(*t*) = *c*_*l*_(1 − *t*)^*−*1^, and (ii) given 𝒯_*l*_ and 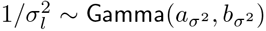, generate triples 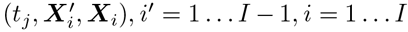 by a scaled Brownian motion upon 𝒯_*l*_. After discarding 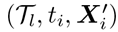, the leaf locations ***X***_*i*_ form an *I* by *J* observed data matrix **X**_*l*_. In Algorithm S1, ABC repeats the procedure above to generate *N* ^syn^ synthetic data (see Figure S2).

**Figure S2:**
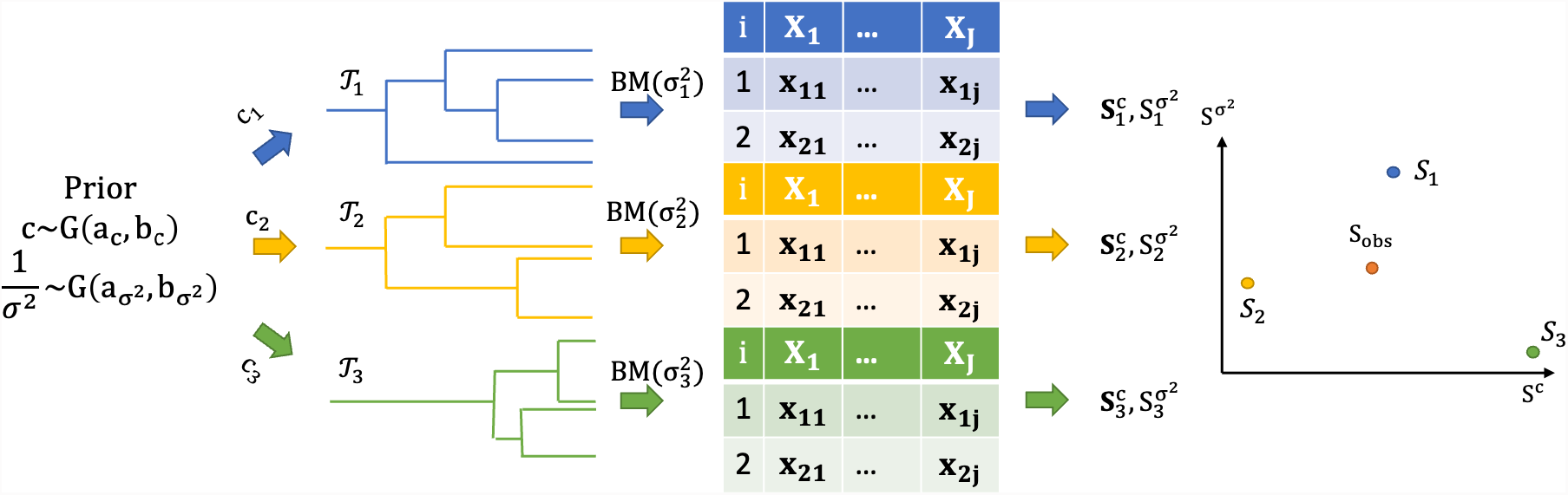
Schematic diagram of synthetic data generation and the calculation of summary statistics (first stage of Algorithm S1). *S*_obs_ is calculated based on the actual observed data.

#### Regression adjustment in ABC

Originally proposed in Beaumont et al. [2002] and later generalized by Blum [2010], regression adjustment for ABC is performed in Step 8 of Algorithm S1. The motivation is to use smoothing technique to weaken the effect of the discrepancy between the summary statistic calculated from synthetic data and that from the observed data. We briefly describe the the procedure of *c*. Additional details can be found in Beaumont et al. [2002] and Blum [2010]. Suppose we are given the observed summary statistics 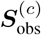 and unadjusted samples 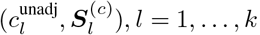, we can calculate the weight for each sample by

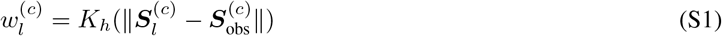

where the bandwidth *h* is set at the largest value, such that 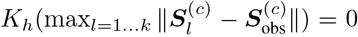 to ensure non-zero importance weight for *k* samples [Sisson et al., 2019] and mean integrated square error consistency [Biau et al., 2015]. Regression adjustment seeks to produce adjusted samples *c*_*l*_ but maintain the sample weights and thus assumes the following model for the unadjusted samples *c*^unadj^ with mean-zero i.i.d errors *ϵ*_*l*_ where 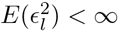 for *l* = 1 …, *k*:

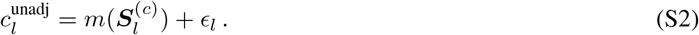

The estimated regression function 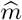 is then a kernel-based local-linear polynomial obtained as a solution of argmin 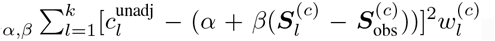. Using the empirical residuals 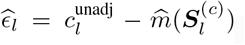, we then construct the adjusted values 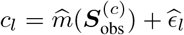.

#### Posterior summary of Euclidean parameters (*c, σ*^2^)

The first stage of our ABC-MH algorithm produces weighted samples 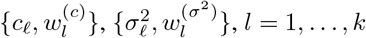, and we summarize the weighted samples as follows. We illustrate the calculations with *c*, and the calculations for *σ*^2^ follow similarly. We calculate the posterior median and 95% credible interval by finding the 50, 2.5 and 97.5% quantiles, and use the posterior median for the second stage of the proposed ABC-MH algorithm when sampling the tree. In general, for calculating the *q* × 100% quantile, we fit an intercept-only quantile regression of *c*_*ℓ*_ with weights 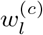; this is implemented by `rq` wrapped in the summary function `summary.abc` in the R package `abc`.

### S2.2 MH Algorithm for Updating the Tree in the DDT Model

In the second stage of Algorithm S1, we have used existing MH tree updates [Knowles and Ghahramani, 2015]. We briefly describe the proposal for generating a candidate tree 𝒯 ′ from the current tree 𝒯 and the acceptance probability. Given the current tree, a candidate tree is proposed in two steps: (i) detaching a subtree from the original tree, and (ii) reattaching the subtree back to the remaining tree (see Figure S3). In Step i, let (𝒮, *ℛ*) be the output of the random detach function that divides the original tree 𝒯 into two parts at the detaching point *u*, where 𝒮 is the detached subtree and *ℛ* is the remaining tree. In this paper, we generate the detaching point *u* by uniformly selecting a node and taking the parent of the node as the detaching point. In Step ii, for the re-attaching point *v*, we follow the divergence and branching behaviors of the generative DDT model by treating subtree 𝒮 as a single datum and adding a new datum 𝒮 to *ℛ*. Given the point *v*, a candidate tree 𝒯 ′ results by re-attaching 𝒮 back to *ℛ* at point *v*. The time of re-attaching point *t*_*v*_ is then earlier than the time of the root of 𝒮 to avoid distortion of 𝒮: *t*_*v*_ *< t*(root(𝒮)). By choosing *u* and *v* as above, we have described the proposal distribution from 𝒯 to 𝒯 ′, *q*(*v*, ℛ), which is essentially the probability of diverging at *v* on the subtree ℛ. The acceptance probability is then

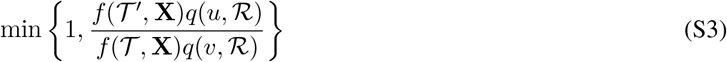

where 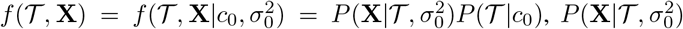 is the likelihood of the tree structure (Proposition 1), *P* (T |*c*_0_) is the prior for the tree (the first two terms in Equation (4)), and *c*_0_ and 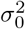 are representative value chosen from the posterior sample of *c* and *σ*^2^, respectively.

## S3 Tree Projection of Pairwise iPCP Matrix

In the Main Paper Section 3.2, we mentioned that a pairwise iPCP matrix **Σ** with entries iPCP_*i,i*′_, *i, i′* = 1, …, *I* need not to be a tree-structured matrix and we address the projection of Σ on to the space of tree-structured matrices here. Given *L >* 1 posterior trees with *I* leaves and the corresponding pairwise iPCP matrix **Σ** = (iPCP_*i,i*′_), each entry of iPCP matrix can be express as 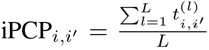, where 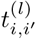, is the divergence time of leaves *i* and *i′* in the *l*-th posterior tree. Obviously, every entry of the iPCP matrix takes the element-wise Monte Carlo average over *L* tree-structured matrix and breaks the inequalities (2) and (3) in the Main Paper. Following the work of Bravo et al. [2009], by representing a tree as a tree-structured matrix, we can project Σ on to the closest tree-structured matrix in terms of Frobenius norm. The projection can be formulated as a constrained mixed-integer programming (MIP) problem:

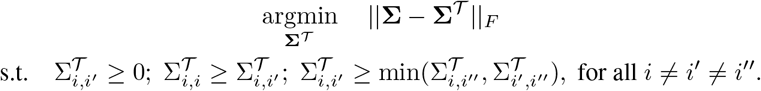

### Algorithm S1 Two-stage hybrid ABC-MH algorithm

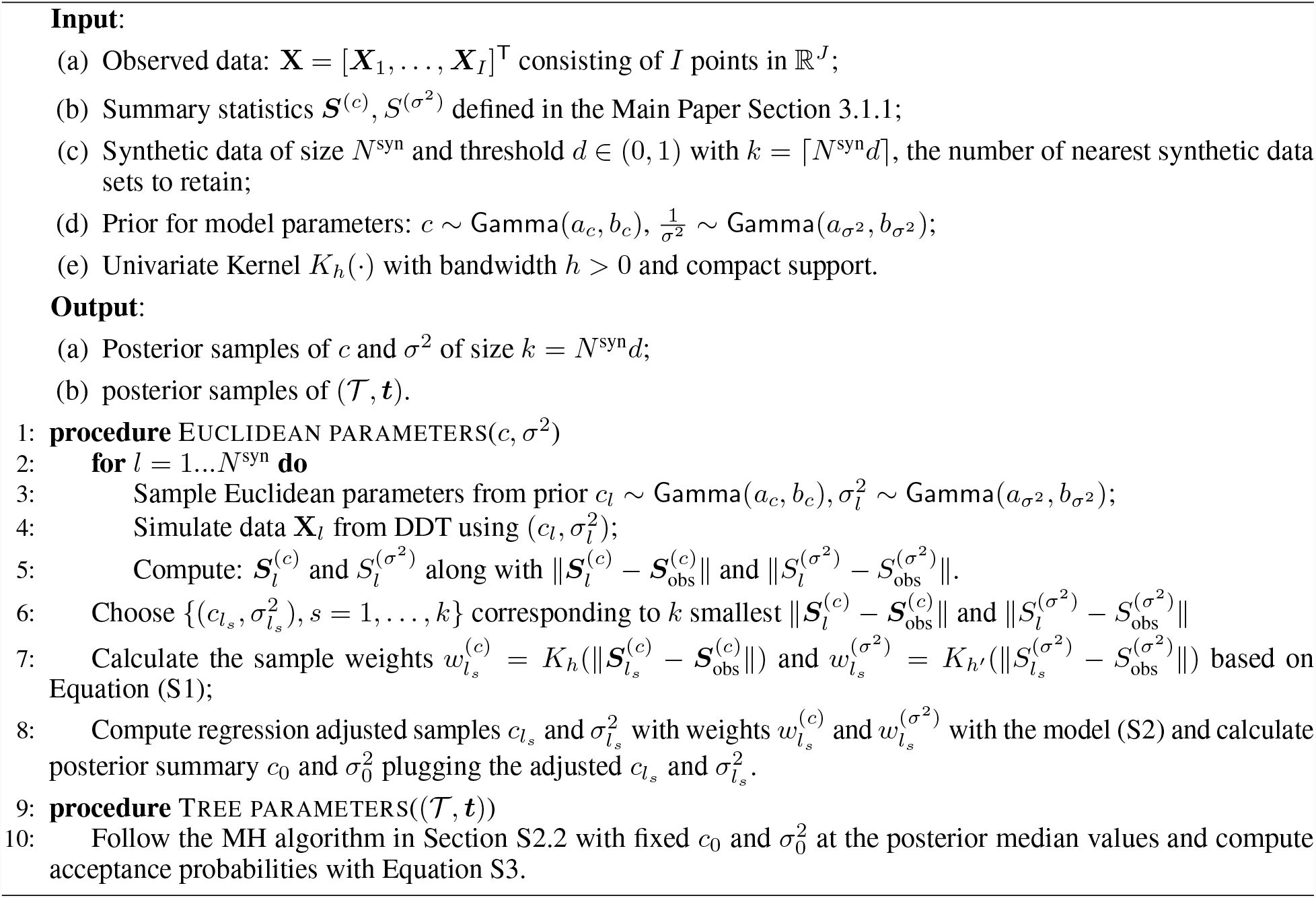

**Figure S3:**
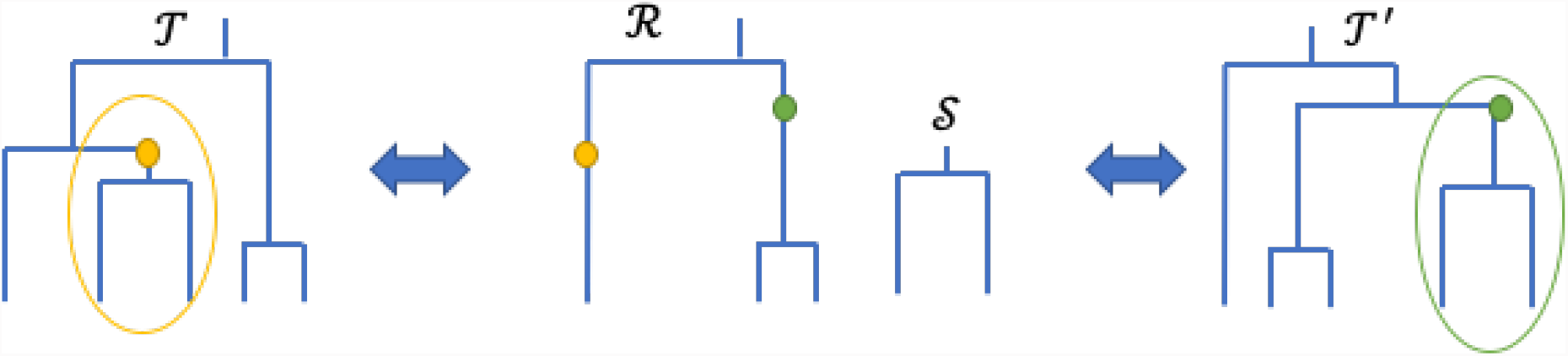
Schematic diagram of proposing a candidate tree in MH. (Left) Current tree 𝒯 with detach point *u* (yellow); (Middle) Intermediate subtrees with remaining tree ℛ and the detached subtree 𝒮; (Right) The proposed tree 𝒯 ′ with reattached point *v* (green).

We applied the projection on the pairwise iPCP matrix from the breast cancer (panel (A)), colorectal cancer (panel (B)) and melanoma (panel (C)) data of NIBR-PDXE and show the result in the Figure S4. In Figure S4, the MAP tree, the tree representation of projected iPCP matrix (MIP tree), the original iPCP matrix and the projected iPCP matrix are shown in from the left to the right columns, respectively. From the left two columns of the tree structures, we found that trees from the MAP and MIP show similar pattern and the MIP tree allows a non-binary tree structure. For example, three combination therapies and two PI3K inhibitors (CLR457 and BKM120) framed by a box form a tight subtree in both MAP and MIP tree, but the subtree in the MIP is non-binary. For the iPCP matrix, high element-wise correlation 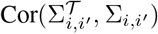 between the original iPCP Σ and the projected iPCP Σ^*𝒯*^ are presented (BRCA: 0.9987; CRC: 0.9962; CM: 0.9918).

**Figure S4:**
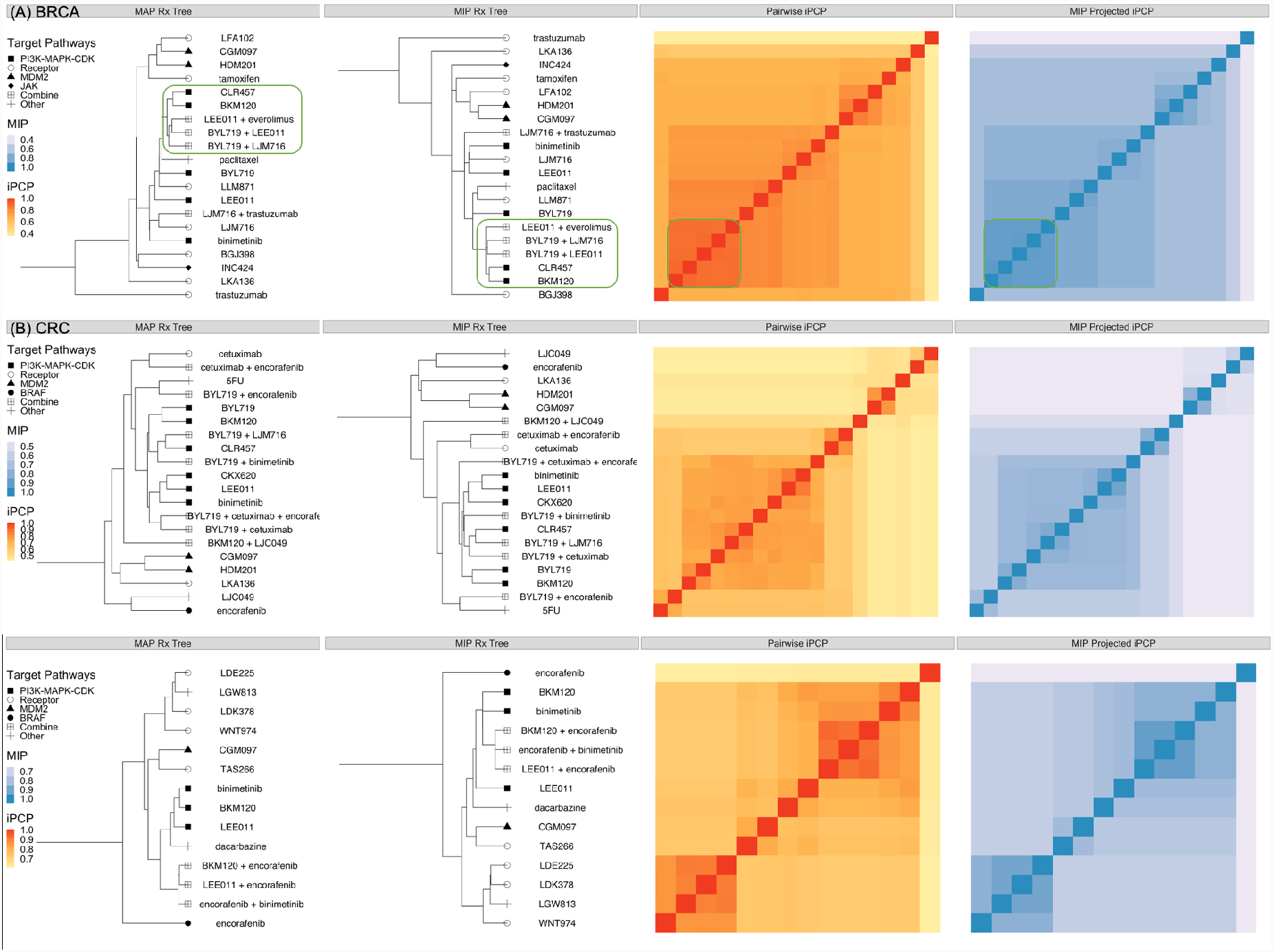
Comparison between (Left two columns) the tree structure from the MAP and the projected iPCP matrix (MIP tree) and (Right two columns) the matrix from the original iPCP matrix and the projected iPCP matrix for (A) breast cancer, (B) colorectal cancer and (C) melanoma. The matrix from the original iPCP and the MIP projected iPCP matrix are aligned by the MIP tree.

## S4 Simulation Studies of Euclidean Parameters

In this section, we empirically compare the Euclidean parameters of *c* and *σ*^2^ from ABC of the proposed two-stage algorithm and single-stage MCMC. We organize this section as follows. We first compare other candidate summary statistics of *c* and *σ*^2^ for ABC in Section S4.1. In Section S4.2, we illustrate the superior inference performance of Euclidean parameters from ABC than single-stage MCMC through a series of simulations. Section S4.3 offers the diagnostic statistics and the sensitivity analysis for ABC stage of the proposed two-stage algorithm and checks the convergence of *c* and *σ*^2^ for the single-stage MCMC.

### Simulation setup

For illustrative purposes, we fixed the observed PDX data matrix with 50 treatments (*I* = 50) and 10 PDX mice (*J* = 10) in all simulation scenarios. In addition, we let *c* and *σ*^2^ take values from {0.3, 0.5, 0.7, 1} and {0.5, 1} respectively to mimic the PDX data with tight and well-separated clusters. For each pair of (*c, σ*^2^), 200 replicated experiments with different tree and observed PDX data matrices were independently drawn according to the DDT generating model. We specify a prior distribution for *c* ∼ Gamma(2, 2) with shape and rate parameterization. For diffusion variance *σ*^2^, let 1*/σ*^2^ ∼ Gamma(1, 1). We compare ABC-MH of the proposed two-stage algorithm against two alternatives based on single-stage MH algorithms [Neal, 2003] (see details in Section S2.2). The first one initializes at the true parameter values and the true tree, referred to as MH_true_. The idealistic initialization at the truth is a best case scenario in applying existing MH algorithm to inferring DDT models. The second alternative, referred to as MH_default_, initializes (*c, σ*^2^) by a random draw from the prior; the unknown tree is initialized by agglomerative hierarchical clustering with Euclidean distance and squared Ward’s linkage [Murtagh and Legendre, 2014] – thus providing a fair apples-to-apples comparison. For the ABC, we generated *N* ^syn^ synthetic data of *c* and *σ*^2^ and kept *k* = ⌈*N* ^syn^*d*⌉ nearest samples in terms of the 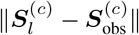 and 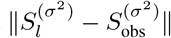. We varied the number of synthetic data *N* ^syn^ and the threshold parameter *d* ∈ (0, 1) under different settings and we specified *N* ^syn^ and *d* in each of the following sections. We ran two MH algorithms with 10,000 iterations and discarded the first 7,000 iterations.

### Performance metrics for Euclidean parameters

We used two algorithm performance metrics to compare our algorithm to the classical single-stage MCMC algorithms. First we computed the effective sample sizes for each Euclidean parameter *c* and *σ*^2^ (ESS_*c*_ and 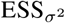) given a nominal sample size (NSS) kept for posterior inference. ESS for each parameter represents the number of independent draws equivalent to NSS posterior draws of correlated (MH_true_ and MH_default_) or independent and unequally weighted samples (ABC stage of the proposed algorithm). We let NSS for MH algorithms be the number of consecutive posterior samples in a single chain after a burn-in period; let NSS for ABC be *k* as in Step 6, Algorithm S1. For *c* and *σ*^2^, the ESS of MH [Gelman et al., 2013] is estimated by 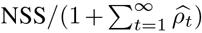 where 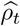 is the estimated autocorrelation function with lag *t* [Geyer, 2011]. *The ESS for ABC [Sisson et al*., *2019] is* the reciprocal of the sum of squared normalized weights, 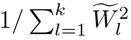, where 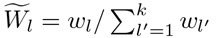 (see weights, *w*_*l*_, in Equation (SS1)). Second, we evaluated how well did the posterior distributions recover the true (*c, σ*^2^). We computed the mean absolute percent bias for *c* and *σ*^2^: | 𝔼 {*c* | **X}** *c −*|*/c* and | 𝔼 {*σ*^2^ | **X}−***σ*^2^ |*/σ*^2^, respectively. We also computed the empirical coverage rates of the nominal 95% credible intervals (CrI) for *c* and *σ*^2^.

#### S4.1 Other Choices of Summary Statistics

Proposition 1 points towards other potential summary statistics for the first stage of Algorithm S1 that uses ABC to produce weighted samples to approximate the posterior distributions for *c* and *σ*^2^. Here we consider a few such alternatives with *N* ^syn^ = 600, 000 and *d* = 0.5% and empirically compare their performances to the summary statistics used in the Main Paper (***S***^(*c*)^ and 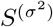) in terms of the mean absolute percent bias in recovering the true parameter values of *c* and *σ*^2^.

### Summary statistic for *c*

Unlike building ***S***^(*c*)^ based on the inter-point distance, the off-diagonal terms of 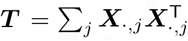 (see the definition of ***T*** in Lemma 1 in Main Paper) is another potential summary statistic for *c*. Since the divergence parameter *c* affects the marginal likelihood implicitly through the divergence time ***t***, the summary statistics for ***t*** is informative for *c*. From Proposition 1, ***T*** is sufficient for *σ*^2^Σ_𝒯_, where the off-diagonal terms of *σ*^2^Σ_𝒯_ taking the form *σ*^2^*t*_*d*_, *d* = 1 … *n* − 1 and containing unrelated information from *σ*^2^. Let ***Q***_***T***_ be a vector of the 10th, 25th, 50th, 75th and 90th percentiles of the off-diagonal terms of ***T***. Because ***T*** is sufficient for *σ*^2^Σ_𝒯_ and involves extra Gaussian diffusion variance parameter, we can design alternative summary statistics based on ***Q***_***T***_ through (i) augmentation, 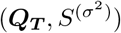 or (ii) scaling, 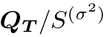. From Figure S5, ***S***^(*c*)^ proposed in the Main Paper outperformed the summary statistics from ***Q***_***T***_ by producing less biased posterior mean estimates.

**Figure S5:**
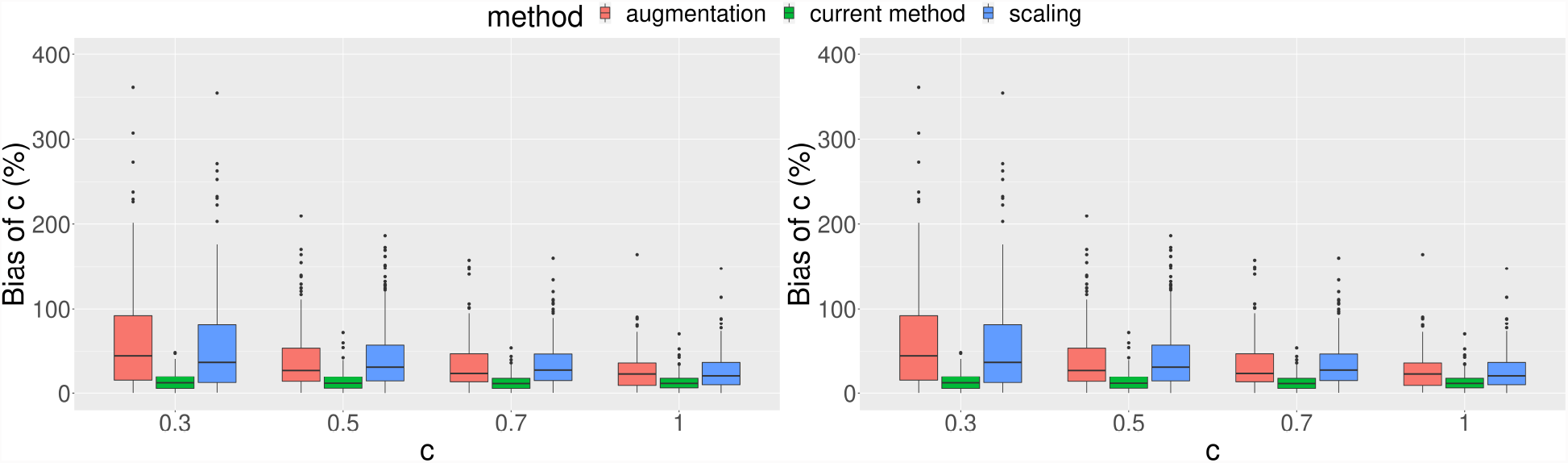
Comparison among different summary statistics for *c* (red: 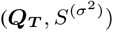; green: ***S***^(*c*)^; blue: 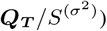) under different values of *σ*^2^ in terms of the mean absolute percent bias. (Left) *σ*^2^ = 0.5; (Right) *σ*^2^ = 1.

### Summary statistic for *σ*^2^

Following Proposition 1, several matrix functionals on the data **X** or statistics ***T*** can be considered as alternatives to 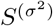. We compare performance of three candidates: (i) average *L*_*1*_ norm (AvgL1) of columns: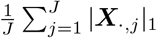; (ii) Frobenius norm of **X**; and, (iii) vector containing 10th, 25th, 50th, 75th and 90^th^ percentiles of first principal component (PC1) of **X**. From Figure S6, the first three methods are comparable while ABC based on principal components shows larger bias due to the information loss.

**Figure S6:**
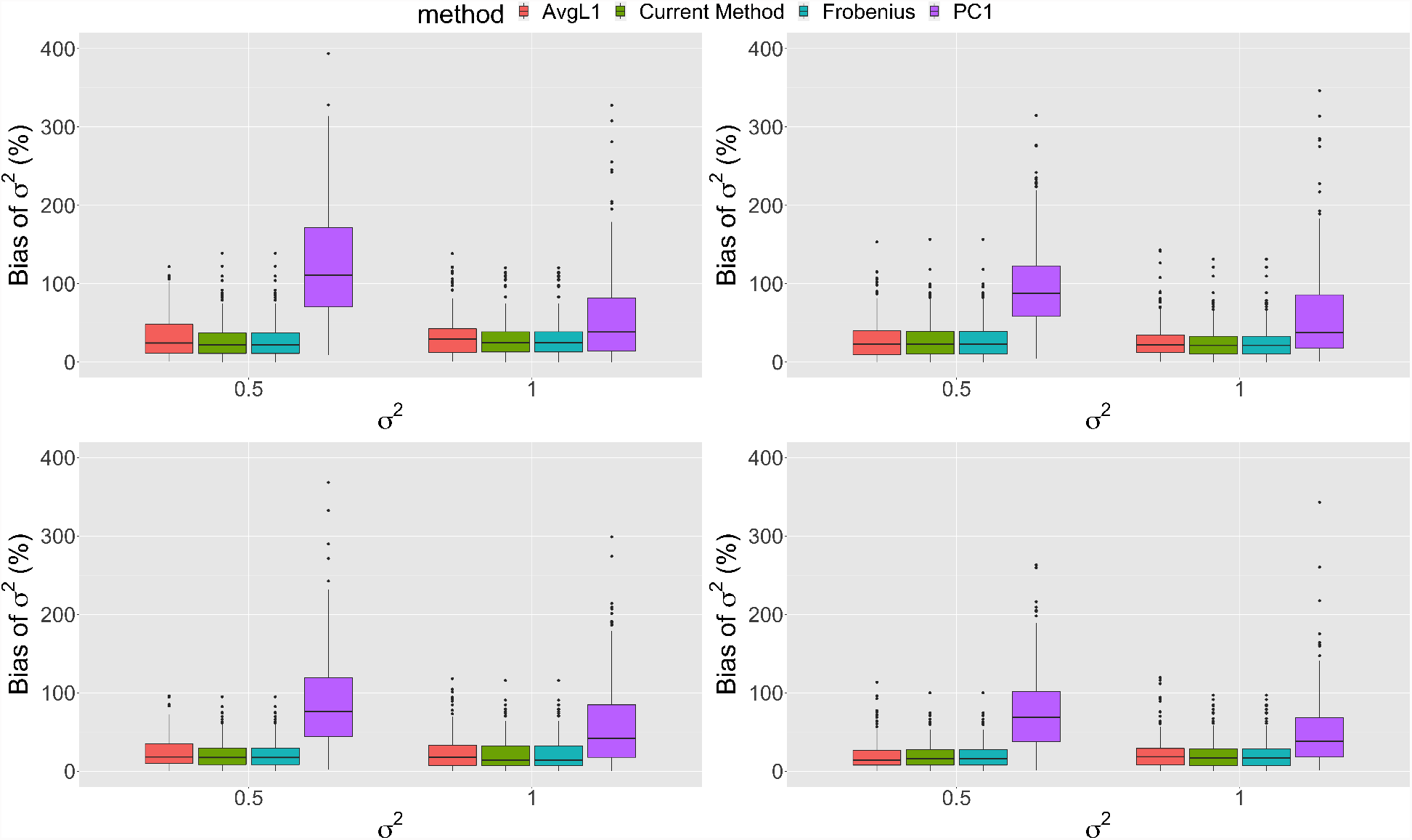
Comparison among different summary statistics for *σ*^2^ under different values of *c* in terms of the mean absolute percent bias. (Upper Left) *c* = 0.3; (Upper Right) *c* = 0.5; (Lower Left) *c* = 0.7; (Upper Right) *c* = 1.0.

#### S4.2 Posterior Inference of Euclidean Parameters

In this section, we show that two-stage algorithm (ABC-MH) outperforms the single-stage MCMC (MH) for real parameters in terms of (i) stable effective sample size (ESS) for (*c, σ*^2^); (ii) similar or better inference on (*c, σ*^2^), as ascertained using mean absolute percent bias and nominal 95% credible intervals.

##### S4.2.1 Stable Effective Sample Sizes of ABC-MH

We calculated ESS-to-NSS ratios at varying truths of *c* and *σ*^2^. To illustrate, we matched the NSS budget of ABC with that of MH (NSS = 3, 000) by keeping *d* = 0.5% of *N* ^syn^ = 600, 000 synthetic data sets that are closest to the observed data in terms of the summary statistic for each parameter (Step 6 of Algorithm S1). Table S1 shows that the ESS_*c*_*/*NSS and 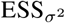/NSS ratio from ABC is stable between 0.64 to 0.68 and around 0.83 across different *c* and *σ*^2^ values, respectively. In contrast, the ESS_*c*_*/*NSS ratio for MH quickly deteriorates (MH_true_: 0.97 to 0.41; MH_default_: 0.73 to 0.35) as *c* increases from 0.3 to 1 and 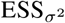/NSS for MH are extremely poor (*<* 0.06) across different values of *c* and *σ*^2^. MH produced very good ESS_*c*_ under small value *c* = 0.3 but poor ESS_*c*_ under *c* = 1. As a result, under larger values of *c*, MH algorithms must run longer to reach a target ESS_*c*_. Although ESS_*c*_ for ABC is not as high as MH_true_ or MH_default_ at *c* = 0.3, the stability of ESS_*c*_ of ABC means that a predictably constant NSS is needed for conducting posterior inference across different values of *c*. Finally, the 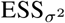 for the diffusion variance parameter from MH algorithms are strikingly smaller than ABC, indicating ABC should be preferred.

##### S4.2.2 Superior Quality Posterior Inference of ABC-MH

Does ABC give better posterior inference with a fixed computational budget? To make fair comparisons, we fixed a total CPU time and used the same computing processor to run the ABC (1st stage of Algorithm 1) and MH algorithms. Let *t*_MH_ and *t*_ABC_ be the estimated CPU time for generating one iteration in MH and one synthetic data in ABC on the same processor. Note, *t*_MH_ includes the additional time for proposing a valid tree. By varying the number of synthetic samples, we can match the total CPU time used by ABC with that of MH algorithms which were run for 10, 000 iterations. We generated 10, 000*t*_MH_*/t*_ABC_ = 17, 345 synthetic data sets and took *d* = 5% with summary statistics ***S***^(*c*)^ and 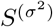 (see different values of *d* in Section S4.3.3) for ABC. Table S2 shows that ABC produced posterior samples that confer comparable inferences about *c* in terms of the bias and coverage of nominal 95% CrIs. The posterior mean of *c* from ABC is comparable to that from MH_true_ and less biased than MH_default_ for all settings. The coverage rates of the nominal 95% CrIs from ABC are comparable to MH_true_ but higher than MH_default_. MH_true_, however, is initialized at true values and is unrealistic in practice. We observed MH_true_ sometimes failed to converge (Table S3), stuck around the initial true values and resulted in deceptively low biases and good coverage rates. Turning to the inference of *σ*^2^, ABC offers a much better alternative to MH algorithms in terms of smaller bias in the posterior mean and better coverage of the 95% credible intervals (Table S2). This is primarily caused by the difficulty of MH in exploring the posterior distribution of *σ*^2^ resulting in chains with high auto-correlations. The squeezed boxplots in Figure S7 indicate that the chains for *σ*^2^ in MH_true_ and MH_default_ were almost always slowly mixing and stuck around the initial values. In addition, unlike the serial nature of MH, ABC can be further parallelized to reduce the wall clock time to a fraction of what is required by MH using multicore processors. Although parallelizing MH with techniques such as consensus MCMC [e.g., Scott et al., 2016] is possible, the parallelized ABC does not require data splitting and will not trade the quality of posterior inference for computational speed.

**Table S1:** ESS-to-NSS ratios between ABC-MH (*d* = 0.5%), MH_true_, and MH_default_. All values here are obtained from 200 independent replications. For each random replication at (*c, σ*^2^). All methods were controlled to produce identical NSS with size 3, 000.

| $c$ | method | ESS/NSS(sd) for $c$ | | ESS/NSS(sd) for $\sigma^2$ | |
| --- | --- | --- | --- | --- | --- |
| | | $\sigma^2 = 0.5$ | $\sigma^2 = 1$ | $\sigma^2 = 0.5$ | $\sigma^2 = 1$ |
| 0.3 | ABC-MH | 0.68(0.032) | 0.67(0.027) | 0.83(0.0048) | 0.83(0.0042) |
| | $\text{MH}_{\text{true}}$ | 0.97(0.11) | 0.96(0.13) | 0.051(0.061) | 0.056(0.072) |
| | $\text{MH}_{\text{default}}$ | 0.73(0.33) | 0.67(0.34) | 0.028(0.043) | 0.038(0.08) |
| 0.5 | ABC-MH | 0.66(0.02) | 0.65(0.018) | 0.83(0.0047) | 0.83(0.0044) |
| | $\text{MH}_{\text{true}}$ | 0.85(0.23) | 0.83(0.24) | 0.034(0.042) | 0.045(0.067) |
| | $\text{MH}_{\text{default}}$ | 0.66(0.35) | 0.62(0.34) | 0.033(0.051) | 0.041(0.067) |
| 0.7 | ABC-MH | 0.65(0.017) | 0.64(0.017) | 0.83(0.0047) | 0.83(0.004) |
| | $\text{MH}_{\text{true}}$ | 0.63(0.31) | 0.67(0.32) | 0.024(0.027) | 0.029(0.038) |
| | $\text{MH}_{\text{default}}$ | 0.53(0.33) | 0.51(0.35) | 0.028(0.039) | 0.038(0.072) |
| 1.0 | ABC-MH | 0.65(0.017) | 0.64(0.017) | 0.83(0.0044) | 0.83(0.0041) |
| | $\text{MH}_{\text{true}}$ | 0.41(0.3) | 0.44(0.32) | 0.019(0.026) | 0.019(0.023) |
| | $\text{MH}_{\text{default}}$ | 0.35(0.29) | 0.35(0.29) | 0.022(0.026) | 0.022(0.027) |

#### S4.3 Algorithm Diagnostics

Here we examine the convergence of MH through the Geweke statistics [Geweke, 1992] and the goodness of fit for ABC. Specifically, two important hyper-parameters are involved in ABC: (i) the kernel bandwidth *h* for samples weights in Equation S1 and (ii) the threshold *d* for *k* = ⌈*N* ^syn^*d*⌉ nearest samples in the Step 6 of Algorithm S1. We follow the test from Prangle et al. [2014] to justify the kernel bandwidth *h* and conduct the sensitivity analysis for threshold *d* to understand how threshold *d* affects the result in terms of the inferential performance.

##### S4.3.1 Convergence of MH Chains in Simulations

In all of our simulations, we ran MH for 10, 000 iterations. Table S3 shows that the percentages of the converged MH chains for 200 replications are between 12.5 and 68.5% within a total 10, 000 iterations (based on Geweke statistic). Running the chains longer will increase these percentages. In contrast, with appropriate choice of bandwidth and the fraction of synthetic samples to keep, ABC does not involve convergence issues and according to Section S4.2 achieves better ESS for a fixed NSS and similar or better quality posterior inference for fixed CPU time.

##### S4.3.2 Diagnostics for ABC

We empirically justify the choice of the kernel bandwidth *h* and the goodness of approximation in ABC algorithm by the calibration method from Prangle et al. [2014] based on the coverage property of the credible interval. Suppose we generated pseudo-observed data **X**_*e*_ in the *e*th replication from the DDT model with parameter 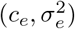, where *c*_*e*_ and 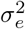 are random draws from the prior 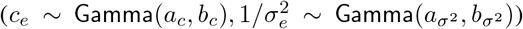 and *e* = 1 … *E*. Once the tuning parameters (*N*^syn^, *d, h*) are decided, Algorithm S1 will output regression adjusted sample 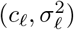 with size *𝓁* = 1, …, *k*; *k* = ⌈*N* ^syn^*d*⌉ based on the input data *D*. We describe diagnostics for *c*, and note that an identical description applies to *σ*^2^ as well. According to Cook et al. [2006], the ABC procedure produces reliable approximations of the posterior if the random variables 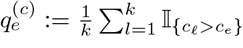 follow a uniform distribution over the interval (0, 1). Accordingly, Prangle et al. [2014] suggest a goodness-of-fit test 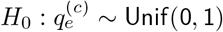 as a diagnostic in order to calibrate ABC. If the test fails to reject the null hypothesis, the empirical quantiles can be viewed as being indistinguishable from the uniform distribution, and the credible interval from the posterior samples would show the asserted coverage. We use the Kolmogorov–Smirnov statistic to carry out the test, follow the simulation setting with *I* = 50 and *J* = 10, and reuse 600, 000 synthetic data sets. The synthetic data is randomly split into two non-overlapping subsets: training data with size 597, 000 and pseudo-observed data with size *E* = 3, 000. Again, we run the ABC part of Algorithm S1 by treating each of the pseudo-observed data sets as the actually observed data with *N* ^syn^ = 597, 000 and *d* = 0.5%. We obtained statistically non-significant KS statistics for *c* and *σ*^2^ (*p*-values: 0.61 for *c*, 0.71 for *σ*^2^). The 95% credible intervals from ABC showed 94.9% and 95.93% empirical coverage rates which are close to the nominal level.

**Table S2:**
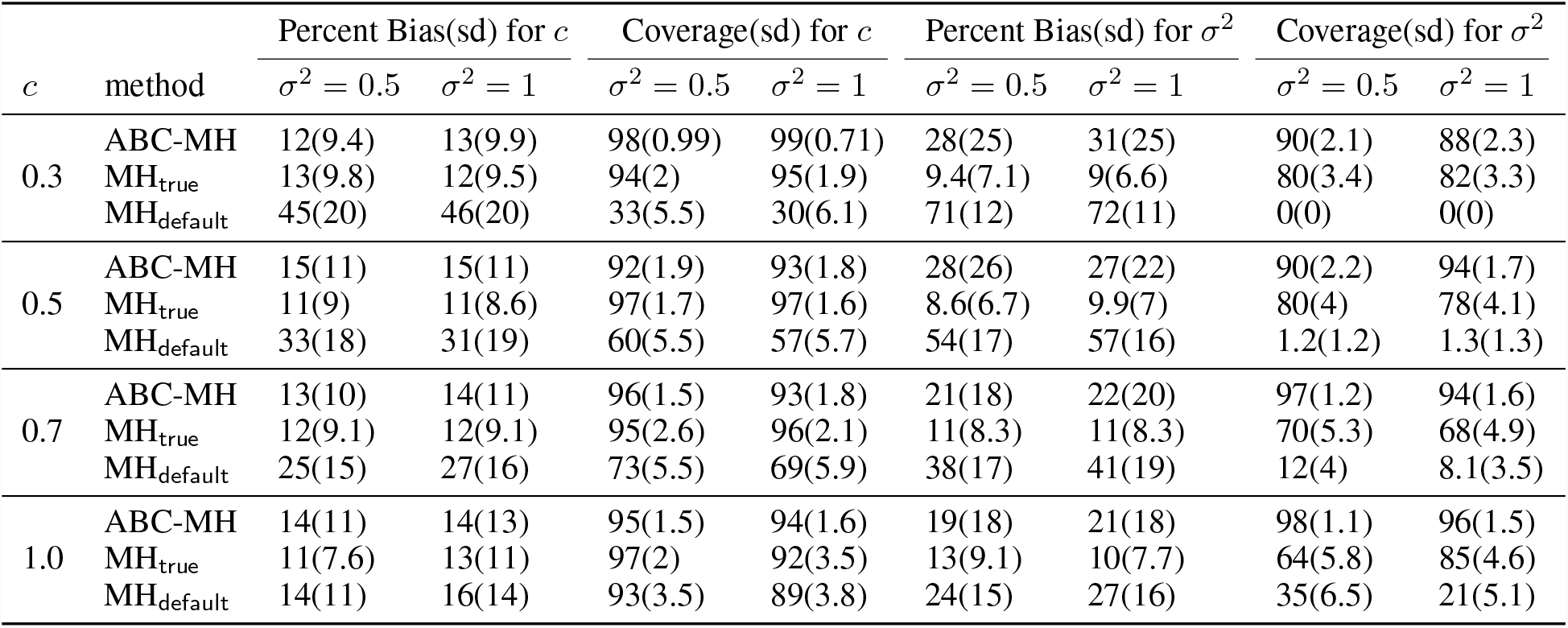
Comparison of inferential performance for *c* and *σ*^2^ between ABC-MH (*d* = 5%), MH_true_, and MH_default_. All values here are obtained from 200 independent replications. For each random replication at (*c, σ*^2^), all methods were run for identical total CPU time and only converged chains from MH algorithms were included.

**Figure S7:**
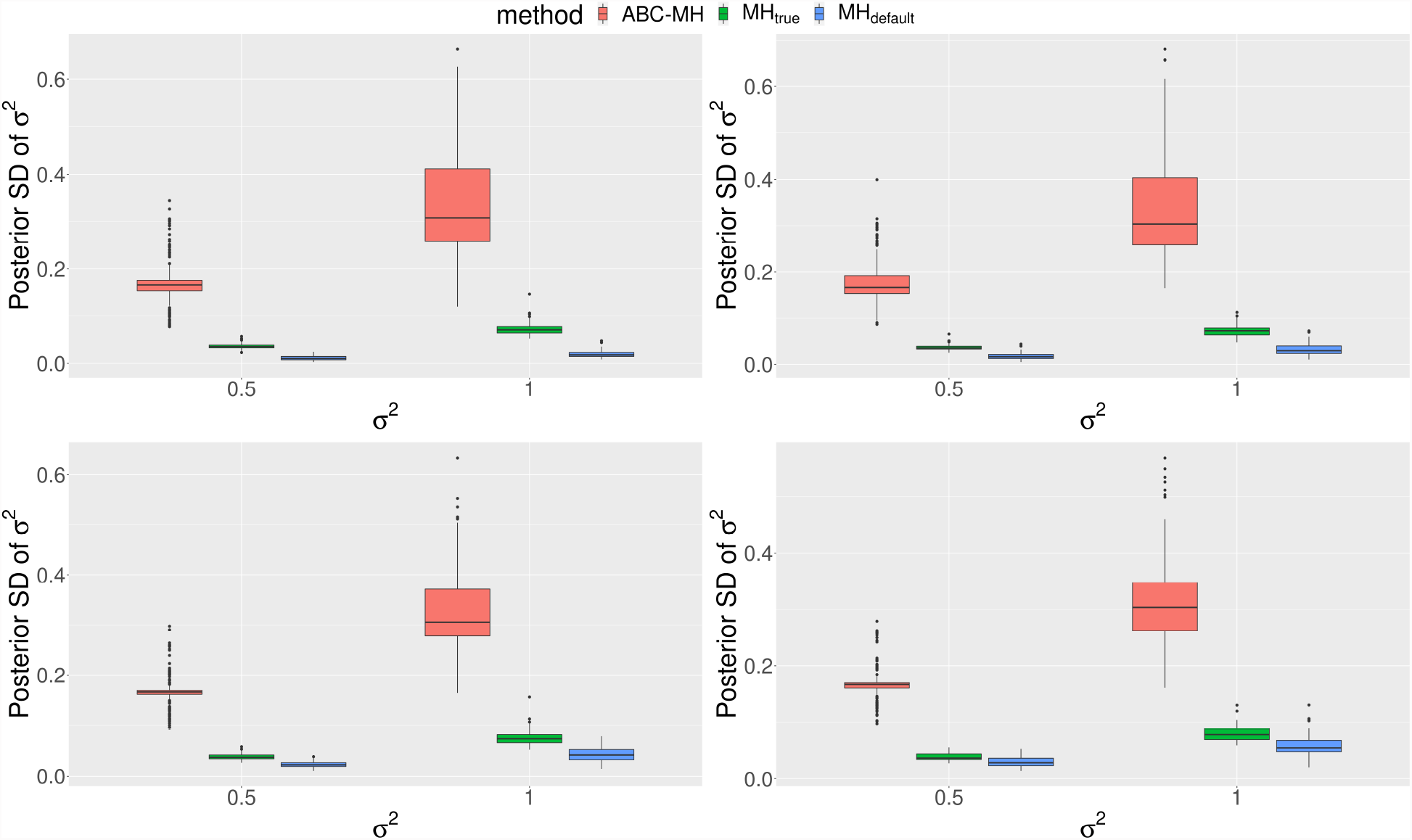
(Upper left) *c* = 0.3; (Upper right) *c* = 0.5; (Lower left) *c* = 0.7; (Lower right) *c* = 1.0. The posterior standard deviation of *σ*^2^ from MH (green and blue) are close to zero across different true *c* showing MH is stuck. Results are based on 200 replications.

**Table S3:** Percentage of converged chains for (i) MH initialized at true (*c, σ*^2^) (MH_true_), and (ii) MH initialized randomly from prior (MH_default_). All values here are obtained from 200 independent replications.

| $c$ | method | Convergence % for $c$ | | Convergence % for $\sigma^2$ | |
| --- | --- | --- | --- | --- | --- |
| | | $\sigma^2 = 0.5$ | $\sigma^2 = 1$ | $\sigma^2 = 0.5$ | $\sigma^2 = 1$ |
| 0.3 | $\text{MH}_{\text{true}}$ | 68.0 | 68.5 | 16.5 | 22.5 |
| | $\text{MH}_{\text{default}}$ | 36.5 | 28.5 | 12.5 | 16.0 |
| 0.5 | $\text{MH}_{\text{true}}$ | 50.0 | 52.0 | 23.0 | 29.5 |
| | $\text{MH}_{\text{default}}$ | 40.5 | 37.5 | 18.5 | 23.5 |
| 0.7 | $\text{MH}_{\text{true}}$ | 38.0 | 46.0 | 26.5 | 27.0 |
| | $\text{MH}_{\text{default}}$ | 33.5 | 31.0 | 14.5 | 25.0 |
| 1.0 | $\text{MH}_{\text{true}}$ | 35.0 | 30.5 | 20.5 | 30.5 |
| | $\text{MH}_{\text{default}}$ | 27.5 | 33.0 | 14.0 | 25.0 |

**Figure S8:**
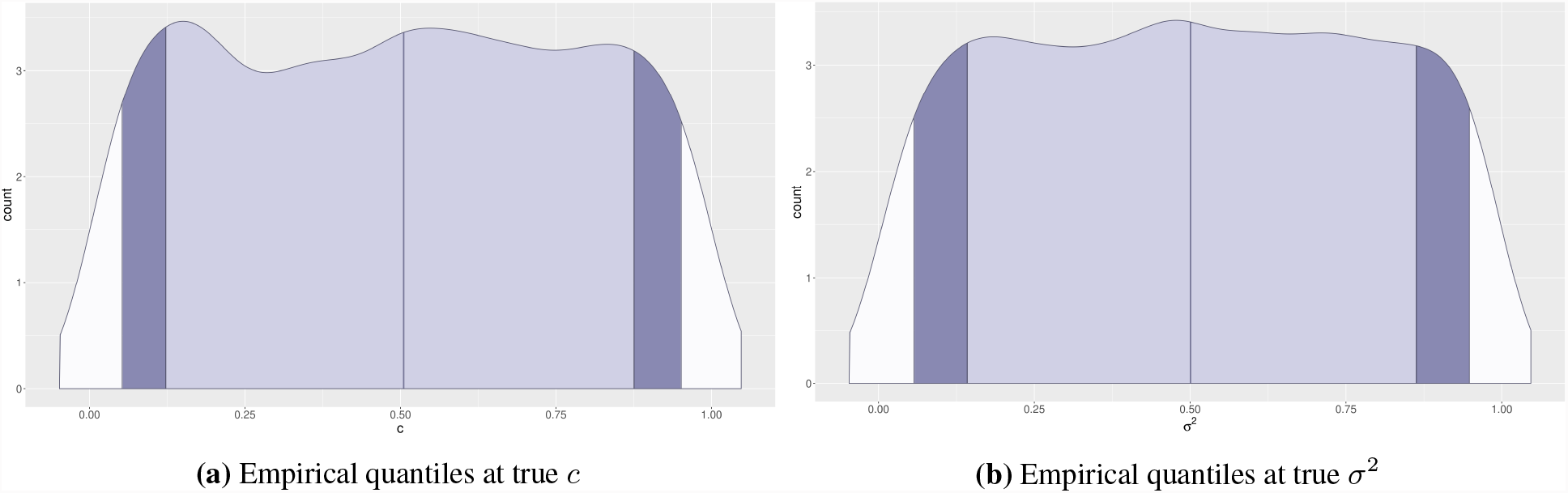
The empirical quantiles at the true value follow the standard uniform distribution indicating calibrated ABC. Results are based on 3, 000 independent draws from the prior.

##### S4.3.3 Sensitivity Analysis of *k* Nearest Samples

In the previous section, we have used a simple diagnostic procedure to show the choice of bandwidth parameter *h* is reasonable. Here we focus on conducting additional simulations to investigate how does varying values of *d* in the Step 6 of Algorithm S1 impact the inferential performance of ABC. We focus on *c* to illustrate the main points. Similar to Table S2 in the Section S4.2 where *d* = 5%, in the following we show the results for *d* = 0.5% and *d* = 1% in Table S4. First, for ABC itself, the bias in the posterior mean is similar, e.g. the mean bias is 14% for all three different *d* when *c* = 1.0 and *σ*^2^ = 0.5. For each pair of (*c, σ*^2^), the empirical coverage rate of the 95% credible interval decreases when *d* increases from 0.5% to 5%. Specifically, the empirical coverage range from 92% to 99% for *d* = 5%, 88% to 97% for *d* = 1% and 84% to 94% for *d* = 0.5%. This is likely caused by a smaller sample size *k* = ⌈*N* ^syn^*d*⌉ and a higher posterior variance under a similar level of bias.

## S5 Additional Simulation Results of R_x_-Trees

In this Section, we provide more simulation results for the Section 4.2 in the Main Paper. We empirically compare the the proposed two-stage ABC-MH with the single-stage MCMC in terms of the MAP tree estimation (Section S5.1) and recovery of pairwise treatment similarities (Section S5.2).

### Simulation setup

For the following simulations, we followed the same setup as in Section S4 with *I* = 50 and *J* = 10, and let *c* and *σ*^2^ take values from {0.3, 0.5, 0.7, 1.0} and {0.5, 1.0 }, respectively. For each pair of (*c, σ*^2^), 50 pairs of tree and data on the leaves were independently drawn based on the DDT model. For ABC, we generated *N* ^syn^ = 600, 000 synthetic data sets from the DDT model with threshold parameter *d* = 0.5%. We assigned priors on *c* ∼ Gamma(2, 2) and 1*/σ*^2^ ∼ Gamma(1, 1) with shape and rate parameterization. We compare the proposed algorithm against two alternatives based on MH algorithms (MH_true_ and MH_default_. We ran MH algorithms (the 2nd stage of the proposed algorithm, MH_true_ and MH_default_) with 10,000 iterations and discarded the first 7,000 iterations.

### Performance metrics

We assess the accuracy of tree estimation using Billera–Holmes– Vogtmann (BHV) distance [Billera et al., 2001] between the true tree and the *maximum a posteriori* (MAP) tree obtained from ABC-MH, MH_true_ and MH_default_, or between the true tree and the dendrogram obtained from hierarchical clustering, respectively. For the pairwise similarities, we follow the Section 4.1 and calculate iPCPs for all pairs of treatments and evaluate the iPCPs by correlation of correlation for estimated similarities and true branching time and the Frobenius norm for the overall matrix.

**Table S4:** Sensitivity analysis of *d* for ABC-MH. We compare the inferential performance for *c* among ABC-MH with *d* = 5%, ABC-MH with *d* = 1%, ABC-MH with *d* = 0.5%, MH_true_, and MH_default_. All values here are obtained from 200 independent replications. For each random replication at (*c, σ*^2^), all methods were run for identical total CPU time and only converged chains from MH algorithms were included.

| $c$ | method | Percent Bias(sd) | | Coverage(sd) | |
| --- | --- | --- | --- | --- | --- |
| | | $\sigma^2 = 0.5$ | $\sigma^2 = 1$ | $\sigma^2 = 0.5$ | $\sigma^2 = 1$ |
| 0.3 | ABC-MH with $d = 5\%$ | 12(9.4) | 13(9.9) | 98(0.99) | 99(0.71) |
| | ABC-MH with $d = 1\%$ | 13(9.8) | 14(10) | 97(1.2) | 96(1.5) |
| | ABC-MH with $d = 0.5\%$ | 14(10) | 15(11) | 94(1.6) | 92(1.9) |
| | $MH_{\text{true}}$ | 13(9.8) | 12(9.5) | 94(2) | 95(1.9) |
| | $MH_{\text{default}}$ | 45(20) | 46(20) | 33(5.5) | 30(6.1) |
| 0.5 | ABC-MH with $d = 5\%$ | 15(11) | 15(11) | 92(1.9) | 93(1.8) |
| | ABC-MH with $d = 1\%$ | 15(12) | 16(12) | 88(2.3) | 90(2.1) |
| | ABC-MH with $d = 0.5\%$ | 16(12) | 16(12) | 84(2.6) | 86(2.4) |
| | $MH_{\text{true}}$ | 11(9) | 11(8.6) | 97(1.7) | 97(1.6) |
| | $MH_{\text{default}}$ | 33(18) | 31(19) | 60(5.5) | 57(5.7) |
| 0.7 | ABC-MH with $d = 5\%$ | 13(10) | 14(11) | 96(1.5) | 93(1.8) |
| | ABC-MH with $d = 1\%$ | 13(10) | 13(11) | 94(1.7) | 90(2.1) |
| | ABC-MH with $d = 0.5\%$ | 13(11) | 14(11) | 90(2.1) | 89(2.2) |
| | $MH_{\text{true}}$ | 12(9.1) | 12(9.1) | 95(2.6) | 96(2.1) |
| | $MH_{\text{default}}$ | 25(15) | 27(16) | 73(5.5) | 69(5.9) |
| 1.0 | ABC-MH with $d = 5\%$ | 14(11) | 14(13) | 95(1.5) | 94(1.6) |
| | ABC-MH with $d = 1\%$ | 14(10) | 14(13) | 88(2.3) | 92(1.9) |
| | ABC-MH with $d = 0.5\%$ | 14(11) | 15(13) | 86(2.4) | 86(2.4) |
| | $MH_{\text{true}}$ | 11(7.6) | 13(11) | 97(2) | 92(3.5) |
| | $MH_{\text{default}}$ | 14(11) | 16(14) | 93(3.5) | 89(3.8) |

#### S5.1 Recovery of the True Tree

The proposed two-stage algorithm decoupled the real and tree parameters, produced better inference for Euclidean parameters (See Section S4.2), resulting in better inference for the unknown treatment tree. In particular, Figure S9 shows that, in terms of the BHV distance, the MAP tree estimates from ABC-MH better recovers the trees than MH_default_ and hierarchical clustering with Euclidean distance and squared Ward linkage (Hclust). On average, MAP from MH_true_ is the closest to the true underlying tree. However, MH_true_ requires knowledge about the truth and is unrealistic in practice. In addition, we observed that the chains from MH_true_ in fact did not mix well and were stuck at the initial values hence falsely appearing accurate. The second stage MH for sampling the tree built on the high-quality posterior samples of *c* and *σ*^2^ obtained from the 1st stage ABC and produced better MAP tree estimates that are on average closer to the simulation truths than MH_default_ and Hclust.

**Figure S9:**
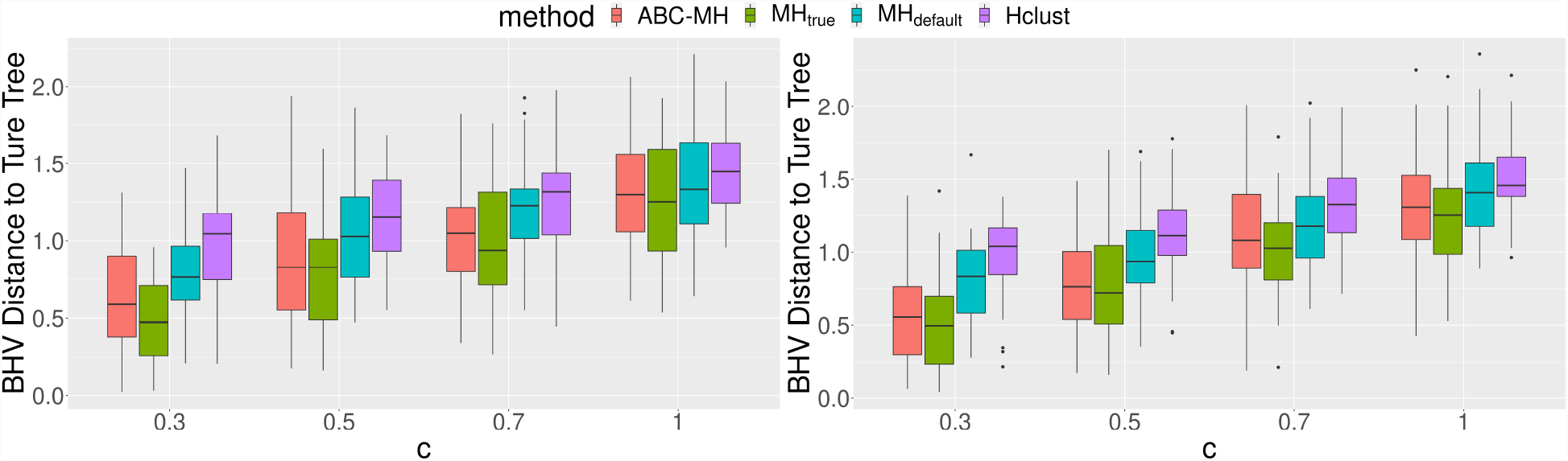
(Left) *σ*^2^ = 0.5; (Right) *σ*^2^ = 1. The BHV distance between the MAP estimate and the underlying tree for each algorithm. Results are based on 50 replications.

#### S5.2 Estimation of Treatment Similarities

The two-stage algorithm also produces better iPCPs due to decoupling strategy and superior inference for Euclidean parameters in the first stage. Similar to the results for MAP, pairwise iPCPs from ABC-MH better recover the true branching time than MH_default_, Hclust and Pearson correlation and reach similar quality to the iPCPs from MH_true_ (See Figure S10). Since MH_true_ requires unrealistic true parameters, MH_true_ is not attainable. From the simulations above, MAP and iPCPs from ABC-MH outperform MH_default_ and take care of overall and local tree details, respectively. We apply the ABC-MH to obtain posterior DDT samples for the real data analysis section.

**Figure S10:**
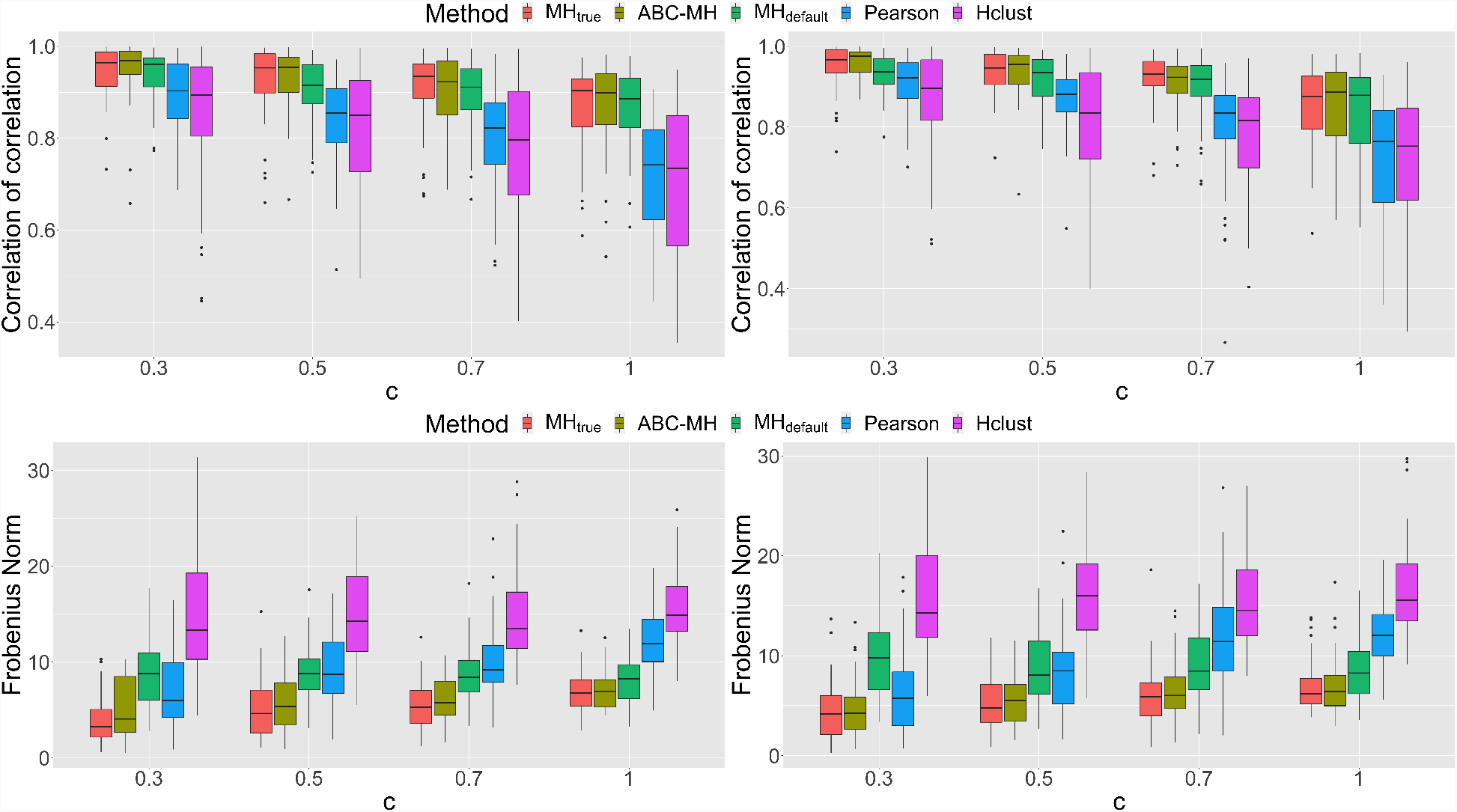
Under different *c* and *σ*^2^, two-stage algorithm better estimates the pairwise similarities than classical single-stage MCMC in terms of correlation of correlation (upper panels) and Frobenius norm (lower panels). (Left) *σ*^2^ = 0.5; (Right) *σ*^2^ = 1. Results are based on 50 replications.

## S6 Additional Results for PDX Analysis

In this section, we provide the pre-processing procedures of NIBR-PDXE and present the results for non-small lung cancer (NSCLC) and pancreatic ductal adenocarcinoma (PDAC) with tables including treatment and pathway information.

### S6.1 PDX Data Pre-Processing

We followed pre-processing procedure in Rashid et al. [2020] and imputed the missing data by k-nearest neighbor method. We take the best average response (BAR) as the response and scale the BAR by the standard deviation over all patients, treatments and across five cancers. Since the scaled BAR contains missing values, we impute the missing data by the k-nearest neighbor with *k* = 10 and compare all treatments to the untreated group. Specifically, we take *x*_*ij*_ = BAR_*ij*_ − BAR_0*j*_, *i* = 1 … *I, j* = 1 … *J* as the observed data, where BAR_0,*j*_ is the untreated BAR for patient *j*.

### S6.2 Test for Multivariate Normal Assumption

From the Proposition 1 and the Equation (5), the observed PDX data follow a multivariate normal distribution. We plot the QQ-plot to check the normal assumption for each cancer and show the result in Figure S11. Among fiver cancers, BRCA (panel (A)) and CM (panel (B)) roughly fall on the 45-degree line and PDAC (panel (E)) is slightly away from the 45-degree line with some extreme data. Contrary, CRC (panel (C)) and NSCLC (panel (D)) deviate from the the 45-degree lines indicating departure from normality.

**Figure S11:**
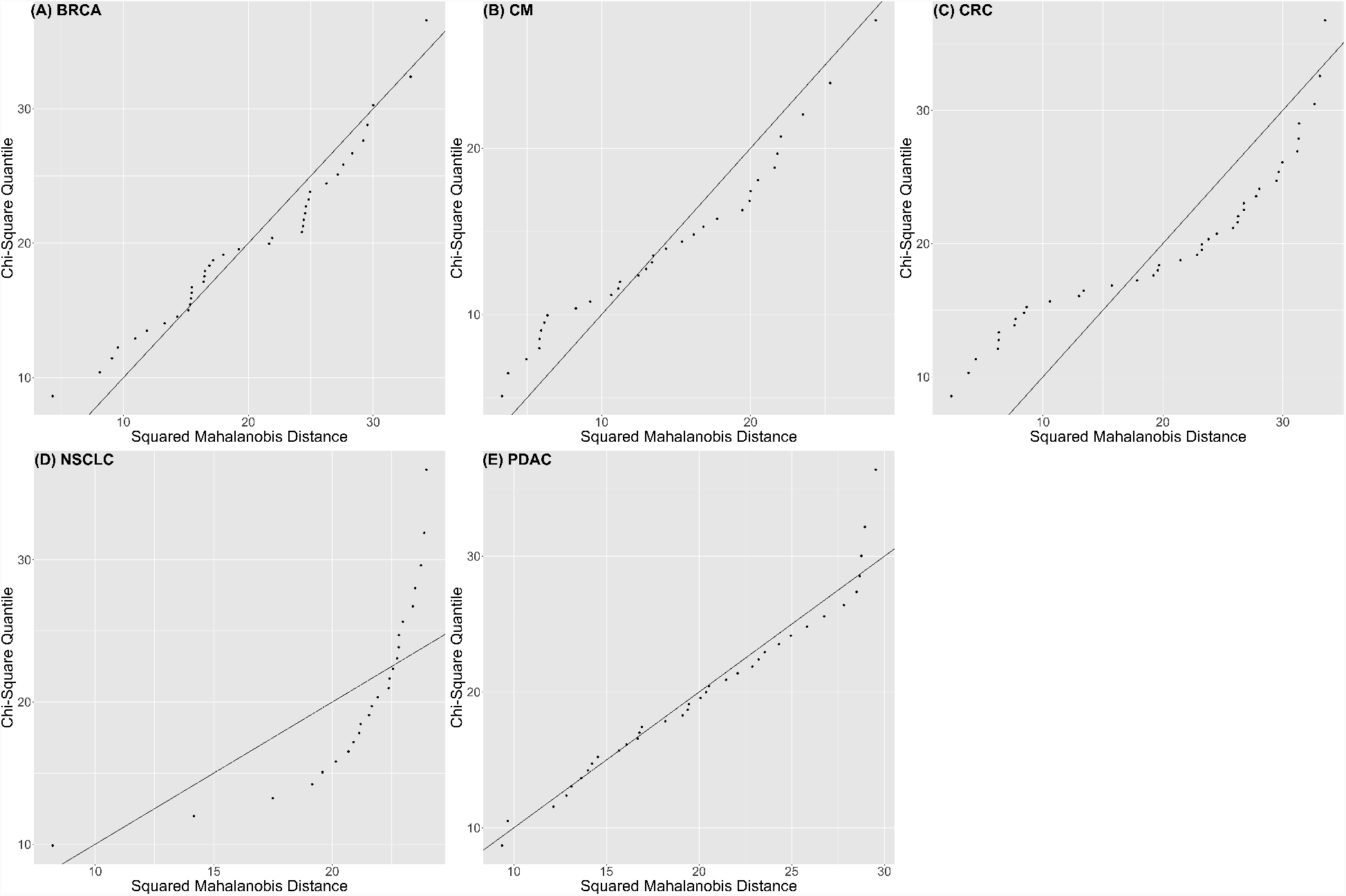
The multivariate normality QQ-plot for (A) breast cancer, (B) melanoma, (C) colorectal cancer, (D) non-small lung cancer and (E) pancreatic adenocarcinoma.

### S6.3 R_x_-Trees for Non-Small Lung Cancer (NSCLC) and Pancreatic Ductal Adenocarcinoma (PDAC)

We applied the R_x_-tree on the rest two cancers in the data: non-small lung cancer (NSCLC) and pancreatic ductal adenocarcinoma (PDAC). Similar to the Figure 5 in the Main Paper, R_x_-tree, pairwise iPCP and (scaled) Pearson correlation are shown in the left, middle and right panels in Figure S12, respectively. Again, we observe that the R_x_-tree and the pairwise iPCP matrix show the similar clustering patterns. For example, three PI3K inhibitors (BKM120, BYL719 and CLR457) and a combination therapy (BKM120 + binimetibin) in NSCLC form a tight subtree and are labeled by a box in the R_x_-tree of Figure S12 and a block with higher values of iPCP among therapies above also shows up in the corresponding iPCP matrix. The R_x_-tree roughly clusters monotherapies targeting oncogenic process (PI3K-MAPK-CDK, MDM2 and JAK) and agrees with the biology mechanism. For example, three PI3K inhibitors (BKM120, BYL719 and CLR457) belong to a tighter subtree in both cancers. Following the same idea as the Main Paper, we further quantify the treatment similarity through iPCP. However, compared to three cancers (BRCA, CRC and CM) in the Main Paper, different problems of model fitting or interpretation lie in NSCLC and PDAC: NSCLC deviates from the normal assumption of Equation (4) (Figure S11) and PDAC shows lower iPCP (average iPCP of PDAC: 0.4119 *<* BRCA: 0.6734, CRC: 0.5653, CM: 0.7535, NSCLC: 0.5817). For concerns raised above, we only verify the model through the monotherapies with known biology for each cancer.

#### Non-small cell lung cancer

Our model suggests high iPCP values for treatments share the same target. For example, our model shows a high iPCP among three PI3K (BKM120, BYL719 and CLR457) inhibitors: (BKM120, BYL719): 0.8402, (BKM120, CLR457): 0.8321, (BYL719, CLR457): 0.8710. For treatments with different targets, our model also exhibits a high iPCP values. For example, the monotherapy HSP990 that inhibits the heat shock protein 90 (HSP90) shows a high iPCP with PI3K inhibitors ((BKM120, HSP990): 0.7108, (BYL719, HSP990): 0.7114, (CLR457,HSP990): 0.7109). Since the inhibiting of HSP90 also suppresses PI3K [Giulino-Roth et al., 2017], it is not surprising to ses a high iPCP between PI3K and HSP90 inhibitors.

**Figure S12:**
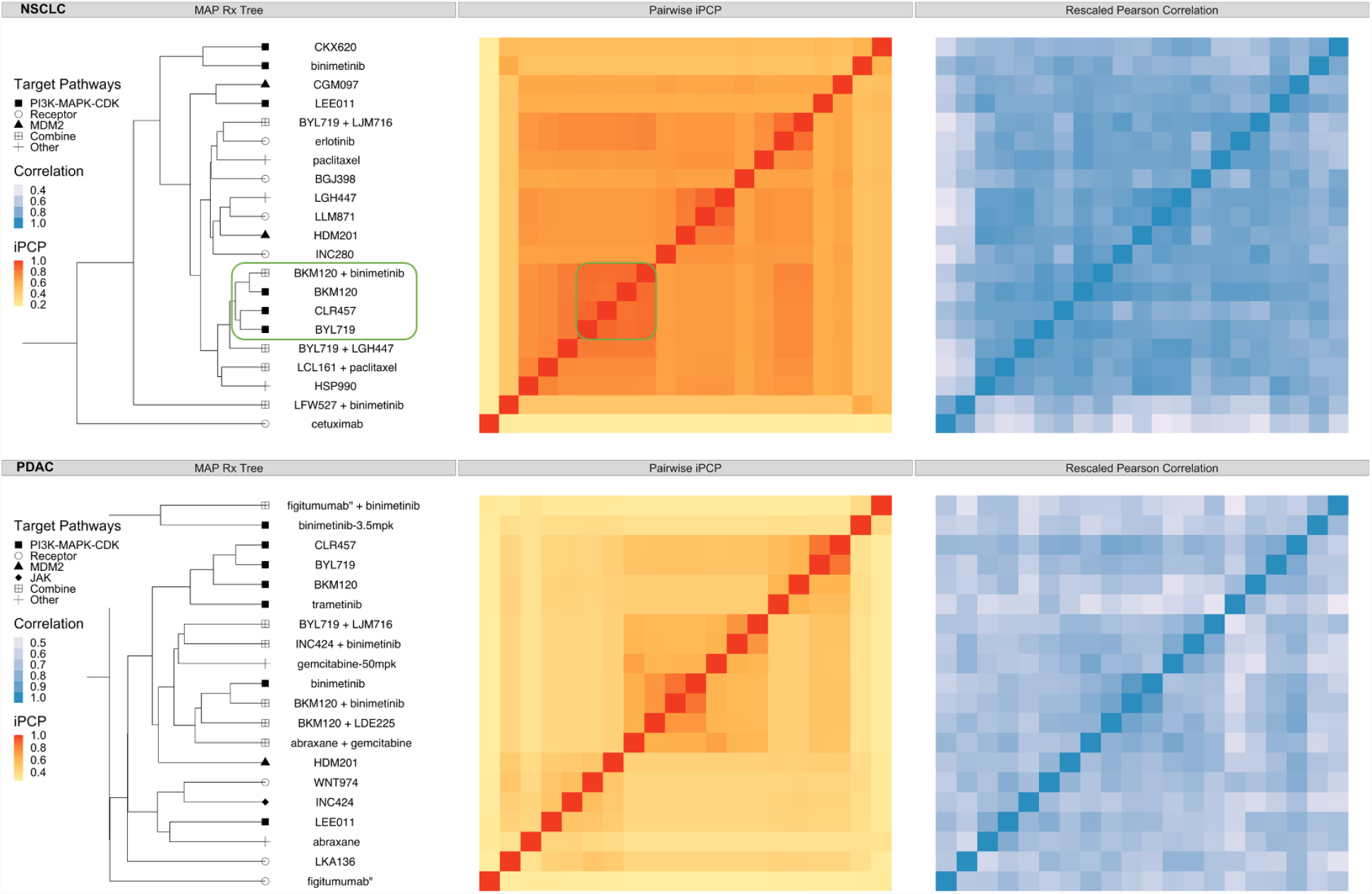
The R_x_-tree and iPCP for non-small cell lung cancer (NSCLC, top row) and pancreatic ductal adenocarcinoma (PDAC, lower row). Three panels in each row represent: (left) estimated R_x_-tree (MAP); distinct external target pathway information is shown in distinct shapes for groups of treatments on the leaves; (middle) Estimated pairwise iPCP, i.e., the posterior mean divergence time for pairs of entities on the leaves (see the result paragraph for definition for any subset of entities); (right) Scaled Pearson correlation for each pair of treatments. The Pearson correlation *ρ* ∈ [−1, 1] was scaled by 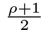 to fall into [0, 1]. Note that the MAP visualizes the hierarchy amongst treatments; the iPCP is not calculated based on the MAP, but based on posterior tree samples (see definition in Main Paper Section 3.2)

#### Pancreatic ductal adenocarcinoma

For PDAC, our model overall suggests a lower iPCP (average iPCP of PDAC: 0.4119). Out of 91 pairs of monotherapies, only BYL719 and CLR457 share a higher iPCP (0.8415). The higher iPCP can be explained by the common target PI3K of BYL719 and CLR457.

### S6.4 R Shiny Application

We illustrate the input and outputs of the proposed method via a R Shiny application hosted on the web (Figure S13). The visualizations are based on completed posterior computations for illustrative purposes. A user needs to specify the following inputs:

A. Cancer type to choose the subset of data for analysis
B. Number of treatments of interest in the subset 𝒜 to evaluate synergy via iPCP
C. Names of the treatments in the subset 𝒜 Given the inputs above, the Shiny app visualizes the outputs:
D. *maximum a posteriori* treatment tree for all the available treatments
E. PCP_*𝒜*_ (*t*) curve for the subset of treatments, 𝒜
F. iPCP_*𝒜*_ value calculated from the corresponding PCP_*𝒜*_ (*t*)

**Figure S13:**
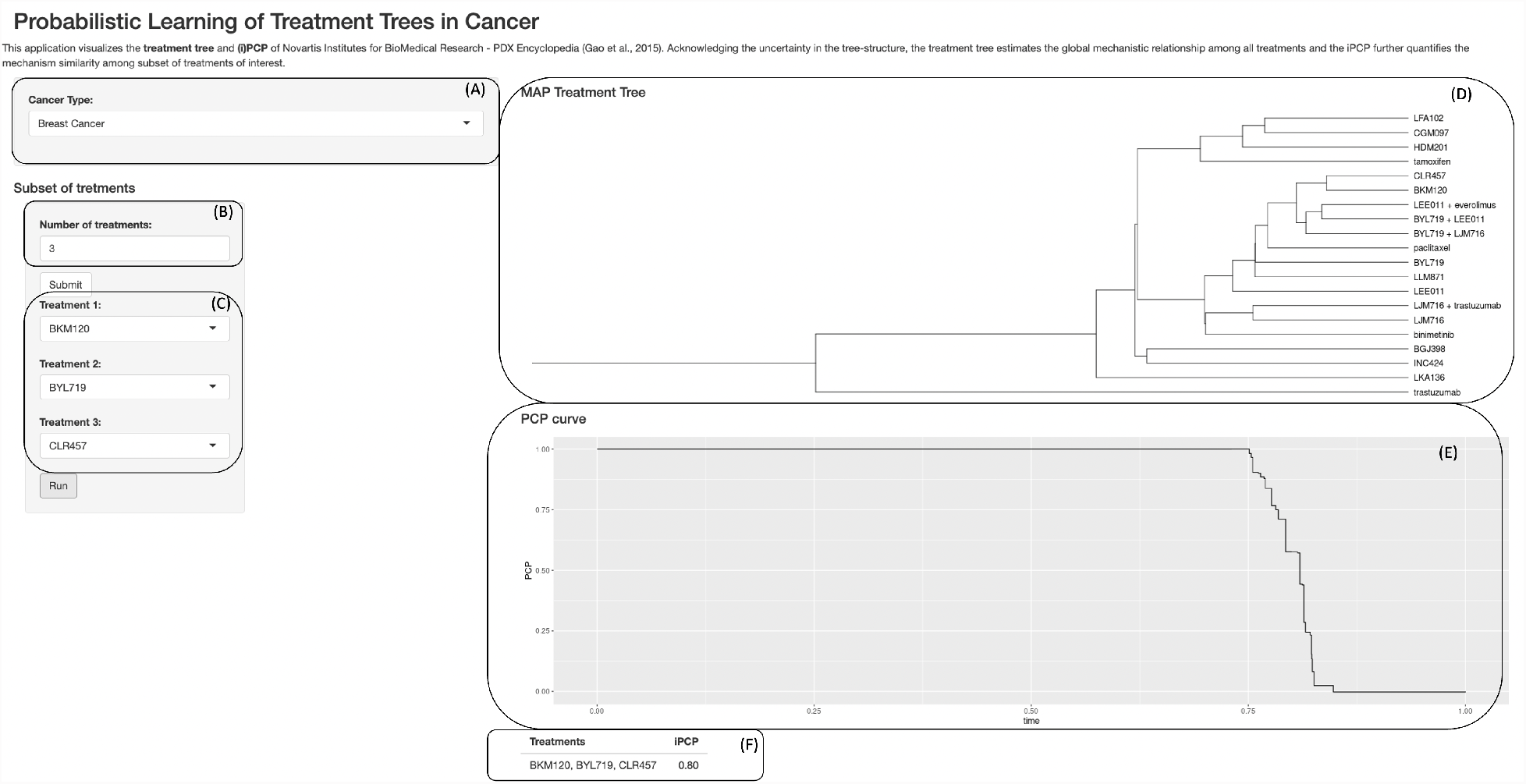
R Shiny app screenshot for illustrating model inputs and outputs for analyzing PDX data (20 treatments for breast cancer); the PCP curve and iPCP value are computed for a subset of three selected treatments.

**Table S5:** Full CPUs series used for computations.

|  |  |
| --- | --- |
| Intel Xeon X series | X5660@2.80GHz |
|  | X5680@3.33GHz |
| Intel Xeon E series | E5-2440@2.40GHz |
|  | E5-2470@2.30GHz |
|  | E5-2450@2.10GHz |
|  | E5-2650v3@2.30GHz |
|  | E5-2650v4@2.20GHz |
|  | E5-2690v4@2.60GHz |
|  | E5-2690v4@2.60GHz |

**Table S6:**
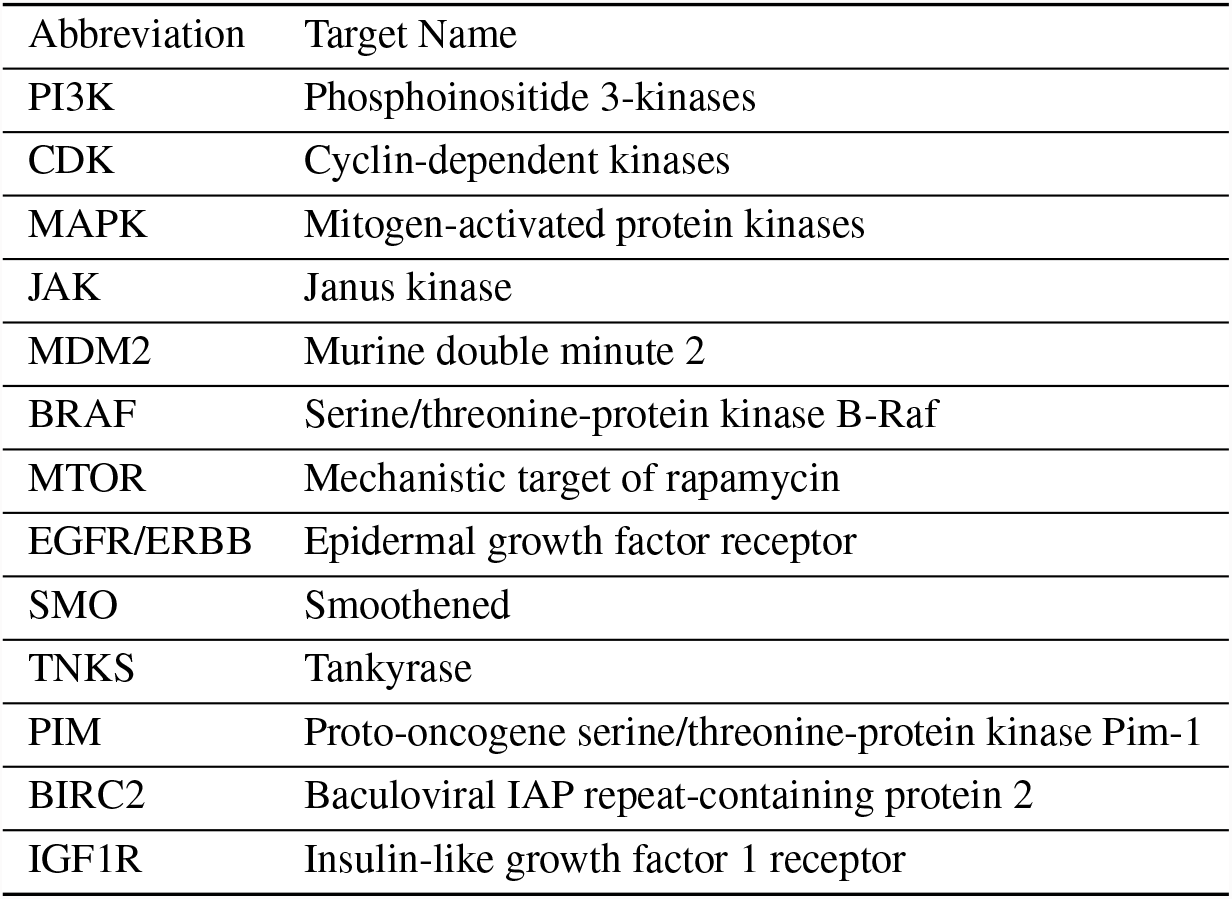
Pathways full names and the corresponding abbreviations.

**Table S7:** Monotherapy names with targets. Different target groups are labeled differently in the Figure 5 and Figure S12.

| Treatment name | Other names | Trade name | Target | Target Group |
| --- | --- | --- | --- | --- |
| 5FU | Fluorouracil | Adrucil | chemotherapy | Other |
| abraxane | nab-paclitaxel | abraxane | Tubulin | Other |
| BGJ398 | Infigratinib |  | FGFR | Receptor |
| binimetinib | MEK162 | Mektovi | MAPK | PI3K-MAPK-CDK |
| BKM120 | Buparlisib |  | PI3K | PI3K-MAPK-CDK |
| BYL719 | Alpelisib | Piqray | PI3K | PI3K-MAPK-CDK |
| cetuximab |  | Erbitux | EGFR | Receptor |
| CGM097 |  |  | MDM2 | MDM2 |
| CKX620 |  |  | MAPK | PI3K-MAPK-CDK |
| CLR457 |  |  | PI3K | PI3K-MAPK-CDK |
| dacarbazine |  | DTIC-Dome | chemotherapy | Other |
| encorafenib | LGX818 | Braftovi | BRAF | BRAF |
| erlotinib | Erlotinib hydrochloride | Tarceva | EGFR | Receptor |
| figitumumab | CP-751871 |  | IGF1R | Receptor |
| gemcitabine |  | Gemzar | chemotherapy | Other |
| HDM201 | Siremadlin |  | MDM2 | MDM2 |
| HSP990 |  |  | HSP90 | Other |
| INC280 | Capmatinib | Tabrecta | MET | Receptor |
| INC424 | Ruxolitinib | Jakafi and Jakavi | JAK | JAK |
| LDE225 | Sonidegib | Odomzo | SMO | Receptor |
| LDK378 | Ceritinib | Zykadia | ALK | Receptor |
| LEE011 | Ribociclib | Kisqal | CDK | PI3K-MAPK-CDK |
| LFA102 |  |  | PRLR | Receptor |
| LGH447 |  |  | PIM | Other |
| LGW813 |  |  | IAP | Other |
| LJC049 |  |  | TNKS | Other |
| LJM716 | Elgemtumab |  | ERBB3 | Receptor |
| LKA136 |  |  | NTRK | Receptor |
| LLM871 |  |  | FGFR2/4 | Receptor |
| paclitaxel |  | Taxol | Tubulin | Other |
| tamoxifen |  | Nolvadex | ESR1 | Receptor |
| TAS266 |  |  | DR5 | Receptor |
| trametinib | GSK1120212 | Mekinist | MAPK | PI3K-MAPK-CDK |
| trastuzumab |  | Herceptin | ERBB2 | Receptor |
| WNT974 |  |  | PORCN | Receptor |

**Table S8:** Combination therapy full names with known targets.

| Combination Therapies | Known Target Pathways | Cancer |
| --- | --- | --- |
| abraxane+gemcitabine | Tubulin+chemotherapy | PDAC |
| BKM120+binimetinib | PI3K+MAPK | NSCLC,PDAC |
| BKM120+encorafenib | PI3K+BRAF | CM |
| BKM120+LDE225 | PI3K+SMO | PDAC |
| BKM120+LJC049 | PI3K+TNKS | CRC |
| BYL719+binimetinib | PI3K+MAPK | CRC |
| BYL719+cetuximab | PI3K+EGFR | CRC |
| BYL719+cetuximab+encorafenib | PI3K+EGFR+BRAF | CRC |
| BYL719+encorafenib | PI3K+BRAF | CRC |
| BYL719+LEE011 | PI3K+CDK | BRCA |
| BYL719+LGH447 | PI3K+PIM | NSCLC |
| BYL719+LJM716 | PI3K+ERBB3 | BRCA,CRC,NSCLC,PDAC |
| cetuximab+encorafenib | EGFR+BRAF | CRC |
| encorafenib+binimetinib | BRAF+MAPK | CM |
| figitumumab+binimetinib | IGF1R+MAPK | PDAC |
| INC424+binimetinib | JAK+MAPK | PDAC |
| LCL161+paclitaxel | BIRC2+Tubulin | NSCLC |
| LEE011+encorafenib | CDK+BRAF | CM |
| LEE011+everolimus | CDK+MTOR | BRCA |
| LFW527+binimetinib | IGF1R+MAPK | NSCLC |
| LJM716+trastuzumab | ERBB3+ERBB2 | BRCA |

